# Prophylactic treatment of *Glycyrrhiza glabra* mitigates COVID-19 pathology through inhibition of pro-inflammatory cytokines in the hamster model and NETosis

**DOI:** 10.1101/2022.05.16.492112

**Authors:** Zaigham Abbas Rizvi, Prabhakar Babele, Srikanth Sadhu, Upasna Madan, Manas Ranjan Tripathy, Sandeep Goswami, Shailendra Mani, Sachin Kumar, Amit Awasthi, Madhu Dikshit

**Author notes:** **Corresponding author** Madhu Dikshit, PhD CSIR-Central Drug Research Institute, Sitapur Rd, Sector 10, Jankipuram Extension, Lucknow, Uttar Pradesh 226031, Amit Awasthi, PhD Immuno-biology Laboratory, Translational Health Science & Technology Institute (THSTI) 3^rd^ Milestone, Faridabad-Gurgoan Expressway, Faridabad, Haryana, India 121001. Equal contribution (Zaigham Abbas Rizvi, Prabhakar Babele). Equal contribution (Srikanth Sadhu, Upasna Madan).

## Abstract

Severe coronavirus disease (COVID-19) is accompanied with acute respiratory distress syndrome & pulmonary pathology, and is presented mostly with inflammatory cytokine release, dysregulated immune response, skewed neutrophil/ lymphocyte ratio, and hypercoagulable state. Though vaccinations have proved effective in reducing the COVID-19 related mortality, the limitation of use of vaccine against immunocompromised, comorbidity, and emerging variants remains a concern. In the current study we investigate for the first-time the efficacy of *Glycyrrhiza glabra* (GG) extract, a potent immunomodulator, against SARS-CoV-2 infection in hamsters. Prophylactic treatment with GG showed protection against loss in body weight and 35-40% decrease in lung viral load along with reduced lung pathology in the hamster model. Remarkably, GG reduced the mRNA expression of pro-inflammatory cytokines and Plasminogen activator inhibito-1 (PAI-1). *In-vitro*, GG acted as potent immunomodulator by reducing Th2 and Th17 differentiation and IL-4 and IL-17A cytokine production. In addition, GG also showed robust potential to suppress ROS, mtROS and NETs generation in a concentration dependent manner in both human polymorphonuclear neutrophils (PMNs) and murine bone marrow derived neutrophils (BMDNs). Taken together, we provide evidence for the protective efficacy of GG against COVID-19 and its putative mechanistic insight, which might be developed as a future immunomodulatory approach against various pathologies with high cytokine production, aberrant neutrophil activation including coronavirus infection.

## Introduction

Severe acute respiratory coronavirus-2 (SARS-CoV-2), the causative agent of the ongoing COVID-19 pandemic, has caused more than 6,228,742 COVID-19 related deaths (https://covid19.who.int/). COVID-19 cases manifest severe pulmonary pathology that is characterized by cytokine release syndrome (CRS), pneumonia, respiratory distress and prothrombotic state (1–3). In addition, SARS-CoV-2 infection also leads to immune dysregulation characterized by lymphopenia, altered neutrophil response, strong pro-inflammatory response and elevated levels of reactive oxygen species (ROS) generation (4). These factors together contribute to pathological symptoms characteristic of COVID-19 not only in the lungs but also in other major organs such as heart, intestine, brain, etc. (5–8). The basis of disease severity is not completely understood, however, factors such as age, dysregulated inflammatory and oxidative stress pathways, and aberrant activation of neutrophils have been implicated. Moreover, a high neutrophil to lymphocyte ratio (NLR) and elevated serum levels of IL-8, the neutrophil chemoattractant cytokine, have been associated with mortality in the COVID-19 patients (9). Sera and postmortem lung biopsies from COVID-19 patients have a high concentration of neutrophil extracellular traps (NETs) components especially in the inflammatory interstitial lesions and airways (10–12). Moreover, activated neutrophils and NETs are known to promote clotting without any other coagulation triggers (11, 13). NETosis is a redox-sensitive phenomenon involving both cytosolic and mitochondrial free radicals. NETs formation is a defensive microbicidal phenomenon to exterminate the invading foreign pathogens but a loss of its control and persistence during inflammation results in the host tissue damage as seen in rheumatic arthritis, diabetes, sepsis, and COVID-19 (10–12).

Though development and active vaccination strategy against COVID-19 have greatly reduced the global morbidity and mortality related to COVID-19, it has been shown that emerging variants of SARS-CoV-2 could not only escape immunity achieved through immunization but also could cause significant morbidity and mortality (14). Furthermore, limited efficacy of vaccine as seen in the case of immunocompromised individuals or individuals with comorbidity remains a future challenge. Strategies to develop anti-viral or immunomodulatory drugs that can inhibit virus growth or mitigate COVID-19 pathologies offers an exciting alternative strategy to counteract COVID-19 related morbidity and mortality (15–17). Pools of evidences now suggest that dietary food and herbal extracts used as prophylactic treatment could mitigate pathogenic infection induced pathologies through anti-viral activities or immunomodulatory effects (18–21). Previously published reports based on computational aided docking or *in-vitro* assays suggest herbal medicines may be used as an alternative remedy either alone or in combination with COVID-19 modern medicines (22). More recently, Jan TS et al 2021 screened 3000 agents from traditional Chinese medicines against COVID-19 by using *in-vitro* assay system and suggested extracts of Ganoderma lucidum, Perilla frutescens, etc to be effective against SARS-CoV-2 (23). However, there are very limited studies which have assessed the efficacy of herbal extracts in the animal models of SARS-CoV-2. Golden Syrian hamsters have been routinely used as a model for COVID-19 as ACE2 receptor of hamsters bear high homology with human ACE2 receptor which results in efficient virus infection and pulmonary pathology upon SARS-CoV-2 infection (24–26).

Here we evaluated the prophylactic efficacy of GG, one of the oldest herbal drugs of Ayurveda (the ancient Indian system of medicine) which is commonly used in Asian and European countries. Different plant parts of GG, especially roots of GG have been used extensively as an alternative or complementary remedy for oxidative and inflammatory diseases because of its several pharmacological effects including anti-viral, anti-microbial, anti-allergic, anti-asthmatic, and immunoregulatory owing to the presence of a myriad of alkaloids, polyphenols, terpenes, flavonoids, coumarins, and other phytochemicals (27, 28). We observed hamsters given prophylactic dosing of GG showed protection in body weight loss, with decrease in lung viral load. Further investigation revealed that GG treatment acted as anti-inflammatory factor and profoundly reduced the expression pathogenic inflammatory cytokines. In addition, GG treatment also suppressed lung injury and lung injury markers most remarkably it led to the inhibition of plasminogen activator inhibitor (PAI-1) which is implicated in COVID-19 induced thrombosis. GG anti-inflammatory activity was found to be through direct inhibition of Th1, Th2 and Th17 cells differentiation *in-vitro*. Our data on human PMNs and mice BMDNs suggests that GG was able to inhibit NETs formation in Telratolimod (TRLM) primed neutrophils after stimulation with PMA or calcium ionophores (A23187 and ionomycin). This effect maybe through GG ability of limiting NETs formation by inhibiting the ROS generation and cytokines release. Taken together, we provide first *in-vivo* report to show that prophylactic treatment with GG protects SARS-CoV-2 infected hamsters and show mechanistically that this protection could be due to direct virus inhibition or through potent immunomodulatory activity.

## Results

### Prophylactic treatment of GG exhibit protective efficacy against SARS-CoV-2 infected hamsters

So far, COVID-19 treatment has mostly relied on active vaccination though a number of potential anti-viral drugs and immunomodulatory compounds have shown to mitigate the clinical pathologies of COVID-19 (14). Remdesivir and Favipiravir have been shown to manage COVID-19 through direct inhibition of viral replication while drugs such as dexamethasone have been shown to act as immune-suppressant thereby reducing the strong inflammatory cytokine release and pulmonary damage (16,17,29). Some, recent studies have pointed that components of GG such as glycyrrhizin, glyasperin A and glycyrrhizic acid could be useful against COVID-19 based on computational docking studies (30–33). Moreover, at least 2 independent groups have shown through in-vitro studies that glycyrrhizin could suppress SARS-CoV-2 pathology through anti-inflammatory or anti-viral properties (30, 31). Based on these reports, we used hamster model for SARS-CoV-2 infection to evaluate the efficacy of GG in mitigating COVID-19 pathologies. To do so, we followed a 5 days pre-treatment regime prior to challenge which was continued until end point i.e. 4 dpi **(Fig. 1A)**. Golden Syrian hamsters receiving GG showed good recovery in body weight as compared to the remdesivir control (R) hamsters (**Fig. 1B**). In line with this, lungs isolated from the euthanized animals on day 4 post challenge showed lower regions of pneumonitis and inflammation as compared to the infected control (**Fig. 2C**). Furthermore, we also found 25-30% decrease in the relative viral load of SARS-CoV2 N2 gene in GG group as compared to the infected group (**Fig. 2D**). Together, our hamster data suggests that prophylactic treatment of GG exhibit protective efficacy against SARS-CoV-2 infection by rescuing the body weight loss and decrease in lung viral load.

**Figure 1:**
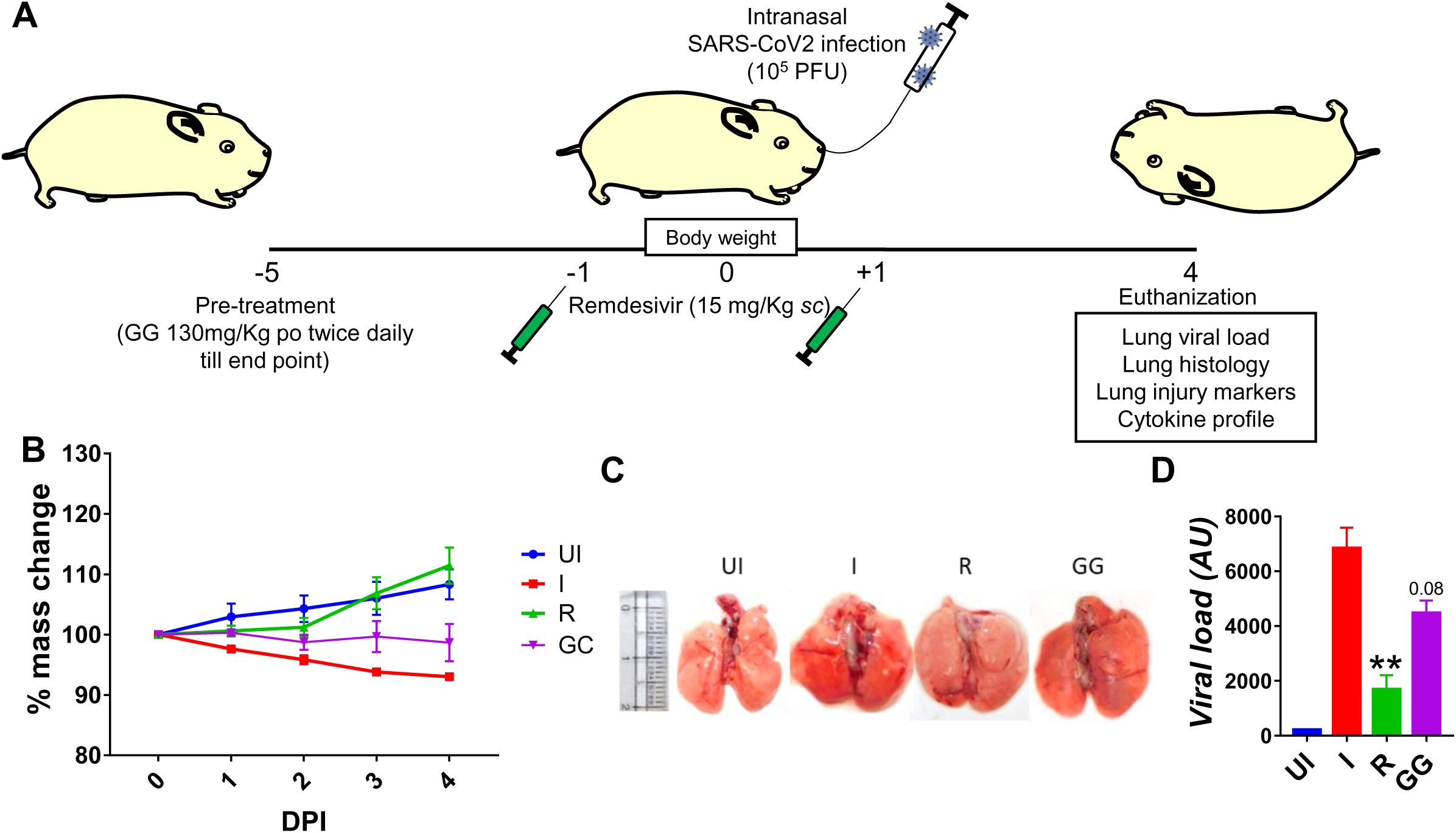
Effect of prophylactic treatment of GG on the SARS-CoV-2 infected hamsters. SARS-CoV-2 infected hamsters were divided into 3 groups viz infection alone (I), infected given remdesivir (R) and infected treated with GG (GG) according to the scheme shown in (A). (B) The changes in body mass were recorded each day post infection and plotted as percent change in the body mass, (C) representative image of the excised lungs showing regions of inflammation and pneumonitis. (D) Relative lung viral load from different groups were estimated by qPCR and shown as bar graph mean ± SEM. N=5 for each experiment. One way-Anova using non-parametric Kruskal-Wallis test for multiple comparison. **P < 0.01.

**Figure 2:**
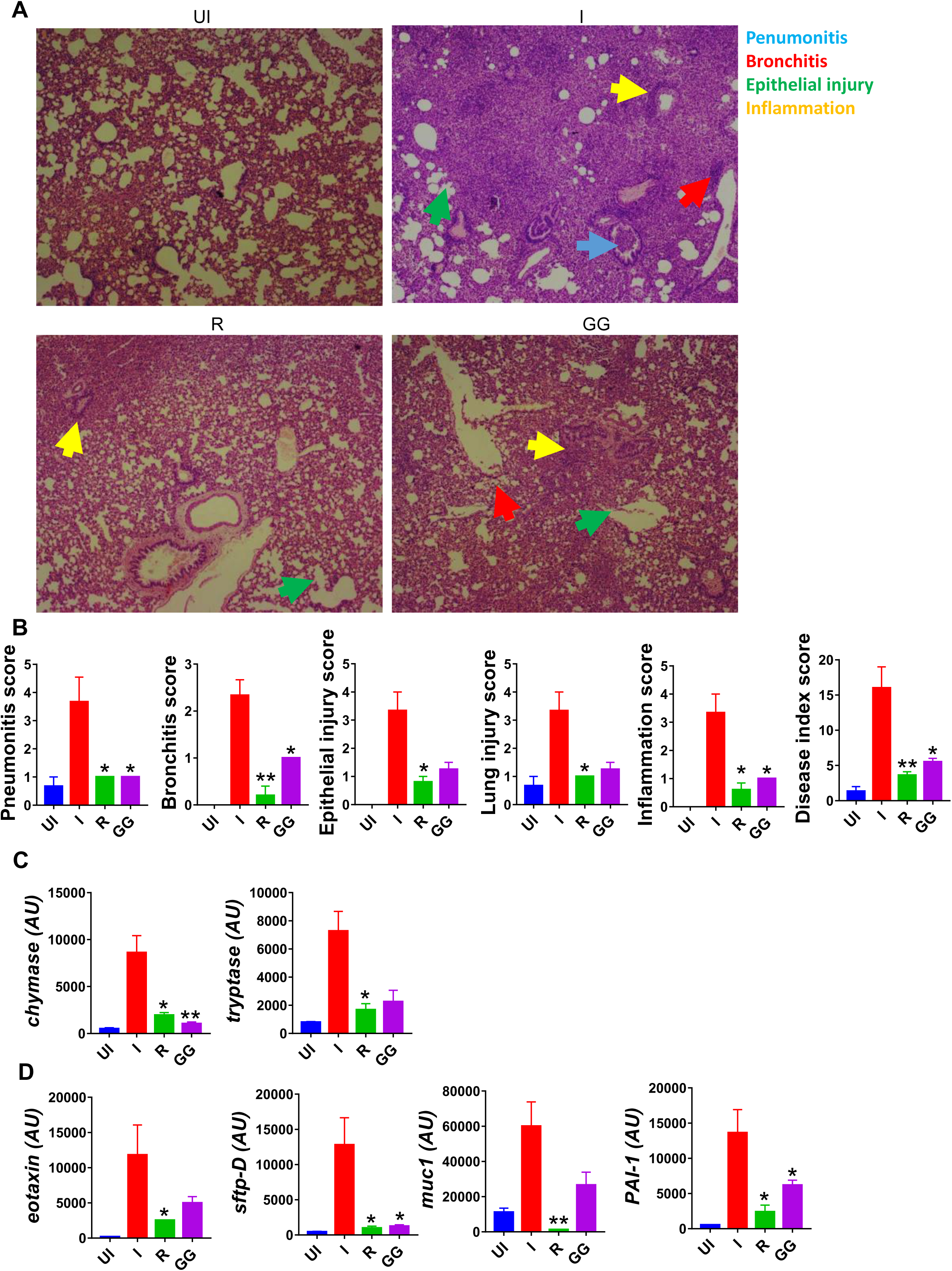
Effect of GG on the lung pathophysiology of SARS-CoV-2 infected hamsters. Left lower lobe of the euthanized hamsters were fixed in formalin, embedded in paraffin, sectioned and stained with H&E. The H&E stained slides were then assessed for histological features and scored. (A) Representative H&E stained lung sections of different groups showing pneumonitis (blue), bronchitis (red), epithelial injury (green) and inflammation (yellow) at 10X magnification. (B) SARS-CoV-2 infected hamsters were divided into 3 groups viz infection alone (I), infected given remdesivir (R) and infected treated with GG (GG) according to the scheme shown in (A). (B) Histological score for pneumonitis, bronchitis, lung injury, epithelial injury and inflammation as assessed by trained pathologist. (C & D) mRNA expression profile of genes associated with mast cell activation or lung injury respectively. N=5 for each experiment. One way-Anova using non-parametric Kruskal-Wallis test for multiple comparison. **P < 0.05, **P < 0.01.

### GG mitigates SARS-CoV-2 induced pulmonary pathology

SARS-CoV-2 infection in clinical cases manifest respiratory distress which is caused by cytokine release syndromes (3). The aggressive release of cytokine in the lungs results in pathological lung injury characterized by pneumonitis, alveolar epithelial injury, etc and elevated levels of injury markers (22, 32). Since prophylactic treatment of GG was able to lower the lung viral, we tried to understand the efficacy of GG to rescue pulmonary pathologies. We therefore carried out detailed histological analysis at 4 dpi of the isolated lung samples. The HE stained lung samples of the GG group showed 2/3-fold reduction in the overall disease index score (comparable to remdesivir control group) with profound mitigation in the pneumonitis, bronchitis, alveolar epithelial injury, lung injury and inflammation histological score as compared to the infection control group (**Fig. 2A & 2B**). One of the important mediators of pathogenic injury of the lung in case of COVID-19 has been identified as mast cells. Our results suggest that GG resulted in suppression of mast cell functional markers such as chymase and tryptase (**Fig. 2C).** Furthermore, we also evaluated the expression of lung injury biomarkers to understand the protection of GG. Prophylactic treatment of GG was found to significantly inhibit the expression of surfactant protein D (sftp-D) and plasminogen activator inhibitor-1 (PAI-1) while reducing the expression of mucin-1 (muc1) and exotoxin to 1.5-2 folds as compared to the infection control (**Fig. 2D).** Taken together, prophylactic treatment of GG showed recovery in the pathological conditions of the infected lungs with lower expression of lung injury markers.

### GG exhibits potent anti-inflammatory property *in-vivo* and limits inflammatory cytokine expression in SARS-CoV-2 infected hamsters

Clinical cases of COVID-19 have been shown to have elevated cytokine profile which known as cytokine respiratory syndrome (3, 34). Heightened levels of pro-inflammatory cytokines lead to tissue injury characteristic of severe cases of COVID-19 (35–37). In line with clinical cases, hamster infected with SARS-CoV-2 shows elevated levels of pro-inflammatory cytokine at 4 dpi as previously reported (24, 38). To understand the immunomodulatory effect of GG on SARS-CoV-2 infected hamsters, we investigated splenomegaly condition and evaluated the cytokine and transcription factor expression in the splenocytes. Spleen of the GG group showed alleviation of hamster’s splenomegaly characteristic of SARS-CoV-2 infection and was comparable to that of the remdesivir group (**Fig. 3A & 3B).** In line with this, mRNA expression profile of pro-inflammatory cytokines such as IFNγ, TNFα, IL4, IL17A and IL13 were all significantly reduced in GG group, however, there was no significant changes observed in the mRNA expression of IL10 and IL6 cytokines (**Fig. 3C & 3D).** In addition, we also evaluated mRNA expression of Th1 and Treg cells viz T-bet and FOXP3 respectively. We did not find any significant changes in the expression of T-bet or FOXP3 in the GG group as compared to the infected control (**Fig. 3E**). Taken together, we show that GG when administered as prophylactic regime lowers the inflammatory cytokine and thus limit the inflammatory cytokine response thereby rescuing the hamsters against COVID-19.

**Figure 3:**
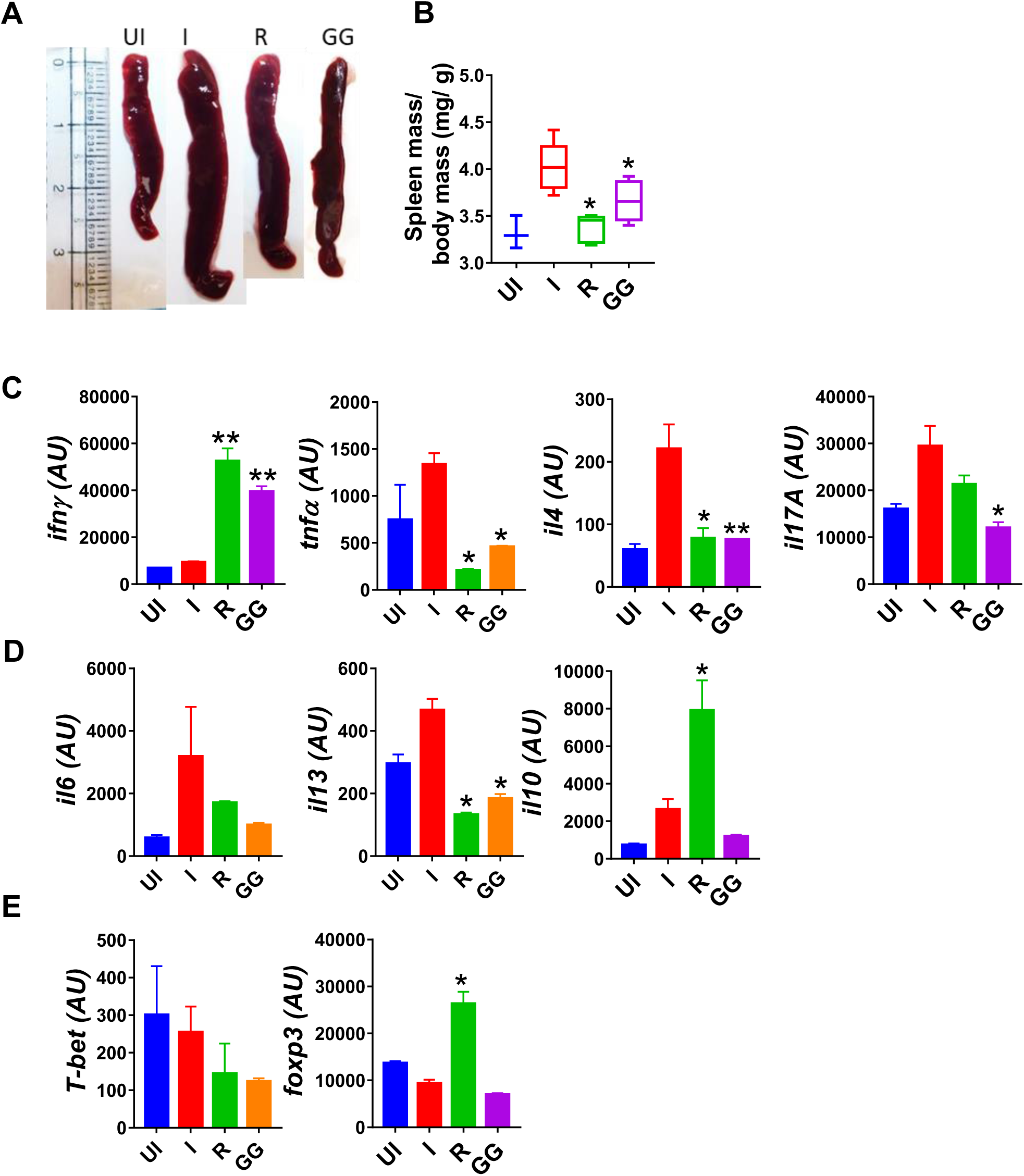
Immunomodulatory effects of GG on SARS-CoV-2 infected hamsters. Immunomodulatory effects of GG were studied in the splenocytes of SARS-CoV-2 infected hamsters and compared with the uninfected control. (A) representative image of excised spleen showing splenomegaly. mRNA expression of key (C & D) inflammatory cytokines and (E) transcription factors. N=5 for each experiment. One way-Anova using non-parametric Kruskal-Wallis test for multiple comparison. **P < 0.05, **P < 0.01.

### GG suppresses Th1, Th2 and Th17 polarization

During acute SARS-CoV-2 infection, T cell-mediated adaptive immune response is required for effective viral clearance and generating long-term antiviral immunity (39). But chronic SARS-CoV-2 infection leads to hyperactivation of T cells and T cell-dependent cytokine release causing immunopathology and poor prognosis (36). Therefore, immune-modulatory drugs are required for treatment of patients with severe lung-inflammation and disease. Dexamethasone (DEX) was used as a positive control since it is a known immunosuppressive drug **(Fig. S1)** (29). It was also recommended for COVID-19 patients with severe respiratory symptoms in RECOVERY trial 2020 (29). In-vitro differentiation of Th1, Th2 and Th17 cells was performed in the presence of graded concentrations of GG and DEX (**Fig. 3 and Fig. S1A-I)**. DEX inhibited the *in vitro* differentiation of Th1, Th2 and Th17 cells with increase in the doses (**Fig. S1**). IC50 value of Dex were calculated and found to be 1.3nM, 3.8nM and 984nM for Th2, Th17 and Th1 cells respectively (**Fig. S1C, F & I**). This showed that Dex is a more potent inhibitor of Th2 cells polarization as compared to Th1 and Th17 cells. To study the immunomodulatory role of GG, we studied its effect on *in-vitro* differentiation of helper T cell subsets Th1, Th2 and Th17 cells **(Fig. 4)**. When incubated with Th2 differentiating cells GG was able to suppress Th2 cells differentiation profoundly even at a dose of 10 µg/ml (**Fig 4A & 4B).** Furthermore, the inhibition of Th2 differentiation was found to be concentration dependent attaining 50% inhibition or IC50 at a concentration of 901.6 µg/ml (**Fig. 4C**). In contrast, inhibition of Th17 cells differentiation was non-significant at lower concentrations with significant inhibition achieved at 300 µg/ml or above concentration (**Fig 4D and 4E**). Furthermore, the inhibition of Th17 differentiation was also found to be concentration dependent reaching 50% inhibition or IC50 at a concentration of 935.4 µg/ml (**Fig. 4F**). Corroborating the above results, we also found profound inhibition of Th1 differentiation in presence of GG at 50 µg/ml and above in a concentration dependent manner (**Fig. 4G & 4H).** The IC50 for GG for Th1 differentiation inhibition was found to be 564.9 µg/ml (**Fig. 4I).** Together, our *in-vitro* data suggests robust immunomodulatory potential of GG to suppress the differentiation of Th1, Th2 and Th17 cells and thereby might contribute to suppress the pathogenic inflammatory response post SARS-CoV-2 infection.

**Figure 4:**
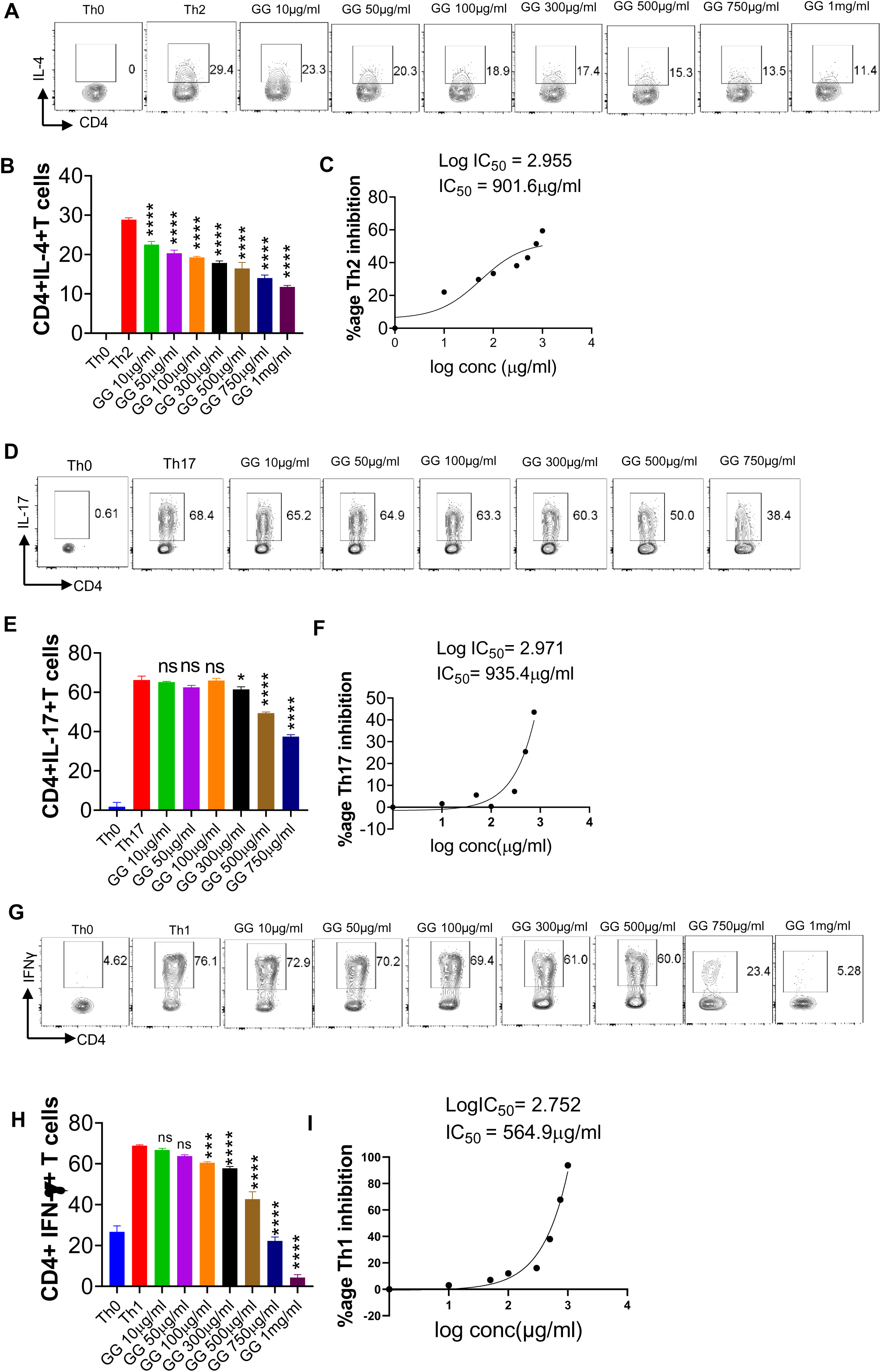
Effect of *Glycyrrhiza gabra* on *in vitro* differentiation of Th1, Th2 and Th17 cells. Spleen and Lymph nodes were isolated from 6-8 weeks old C57BL/6 mice and their single cell suspension was prepared. Cells were activated using soluble anti-CD3 antibody and differentiated into helper T (Th)2 (**A, B**), Th17 cells (**D, E**) and Th1 conditions (**G, H**) using recombinant mouse IL-4; TGF-β + IL-6 and IL-12 cytokines respectively. *Glycyrrhiza gabra* was added in concentrations ranging from 10ug/ml to 1000ug/ml at the start of culture. Cells were differentiated for 72 hours and IL-4, Il-17 and IFNγ production was measured by Intracellular cytokine staining. IC50 values were calculated using Graph pad prism software **(C, F, I)**. *P<0.05, **P<0.01, ***P<0.001, ****P<0.0001 by one-way ANOVA.

### GG exhibited an inhibitory effect on TRLM-PMA/Ionophores stimulated superoxide anions and NETosis in human PMNs and murine BMDNs

Since GG contains various phytoconstituents with myriad of activities, we tested the effect of GG for anti-NETotic activity of neutrophils. Much of the understanding of the mechanism that underlines the NOX-dependent and NOX-independent NETosis resulted from the studies using PMA, A23187, and ionomycin (40, 41). We found comparatively more induction of NETs with A23187 in mice BMDNs and with ionomycin in human PMNs (**Fig. S2A-D**), we therefore used A23187 for mice BMDNs and ionomycin for human PMNs in the subsequent studies. To assess the effect of TLR7/8 receptor activation, neutrophils were pre-treated with TRLM at 10 µM for 30 min, which elicited a significant induction in NETs (51% for PMA, 55% for ionomycin in human and 52% for PMA, 54% for A23187 in mice) (**Fig. 5A-D)**. It is important to note that the concentration of PMA and calcium ionophores used here were submaximal (PMA 12.5 nM; A23187 1.25 µM; ionomycin 2 µg/ml). These results show that TRLM or the two types of inducers alone triggered minimal changes which were increased multi-fold when these were coupled together, indicating the initial priming of TLR7/8 receptors by TRLM and the consequent enhancement of ROS, mtROS, and DNA release by PMA and ionophores.

**Figure 5:**
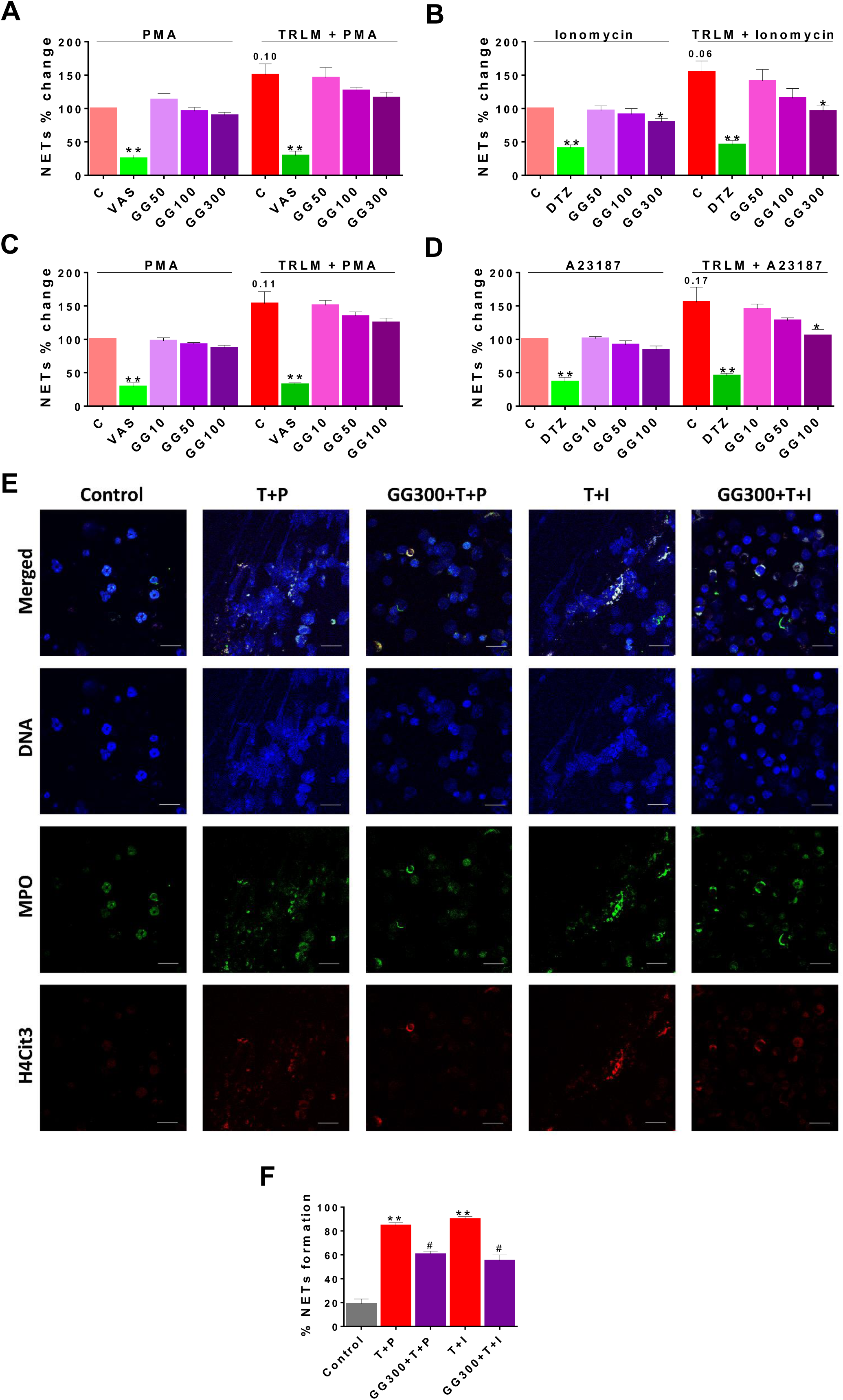
Effect of GG on TRLM primed PMA/calcium ionophores induced cytosolic ROS and mtROS production in human PMNs and murine BMDNs. **(A)** Cytotoxic potential of GG extracts on human PMNs and murine BMDNs. Percent cell death was obtained by flow cytometry using PI (10 µg/ml). Doxorubicin (10 µM) was used as a positive control (100%). **(B-I)** After pre-incubation of PMNs and BMDNs with different concentrations GG, cells were treated with TRLM (10 µg/ml) for 30 min and stimulated with sub-maximal concentration of PMA (12.5 nM) and A23187 (1.25 µM) / ionomycin (2 µM) for 30 min. DCF-DA (10 µM) and MitoSOX (10 µM) were used for cytosolic ROS (**B-C**: PMNs; **D-E**: BMDNs) and mtROS (**F-G**: PMNs; **H-I**: BMDNs) detection, respectively using flow cytometry. A significant increase in ROS and mtROS were induced by TRLM-PMA/ionophores coupling. GG showed a marked inhibitory effect on PMA and ionophore-induced ROS and mtROS production. NAC and MitoTEMPO were used as the positive control for ROS and mtROS assays, respectively. All the data are represented as Mean ± SEM, n = min 3 per group, and statistical analysis consisted of one-way ANOVA followed by Bonferroni’s test (*p<0.05, **p<0.01, ***p<0.001 vs respective control groups; #p<0.05, ##p<0.01 vs PMA/ionophores treated groups). C, control; NAC, N-acetyl cysteine; MT, MitoTEMPO; G10, GG 10 μg/ml; G50, GG 50 μg/ml; G100, GG 100 μg/ml; T, TRLM; P, PMA; I, Ionomycin; A, A23187.

Currently, approaches to prevent NETs generation are considered an attractive strategy to limit uncontrolled inflammation for COVID pathologies (40). Supplementing the cells with GG could inhibit double-stranded DNA release, a hallmark for NETs formation. GG exhibited an inhibitory effect on NETs in a concentration-dependent manner. A low concentration of GG has no significant effect on the inhibition of NETs in neutrophils, however higher concentrations exerted an inhibitory effect on the release of dsDNA (**Fig. 5A & 6B**). Ionomycin-induced NETosis in human PMNs was reduced from 25% to 36% after treatment with 100 and 300 µg/ml, p<0.05 of GG respectively. In contrast, with TRLM-PMA stimulation GG did not exert a noticeable reduction in DNA release; a maximum of 20% inhibition was seen at 300 μg/ml. Treatment of TRLM-A23187 stimulated murine BMDNs with 10, 50, and 100 µg/ml GG resulted in down-regulation of NETosis to 5%, 18%, and 32%, respectively (p<0.05), while exposure to TRLM-PMA resulted in 13% (50 μg/ml) to 22% (100 μg/ml, p<0.05, **Fig. 5C & 6D**). **Fig S3A** show more than 90% cells remains viable up to 300 μg/ml in both human and mice. The IC50 of GG on NETs in BMDNs (100.2 and 89.5 µg/ml using PMA and A23187, respectively) were calculated and all the experiments were conducted using GG concentration based on the calculated IC50 values (**Fig. S3B & S3C**).

**Figure 6:**
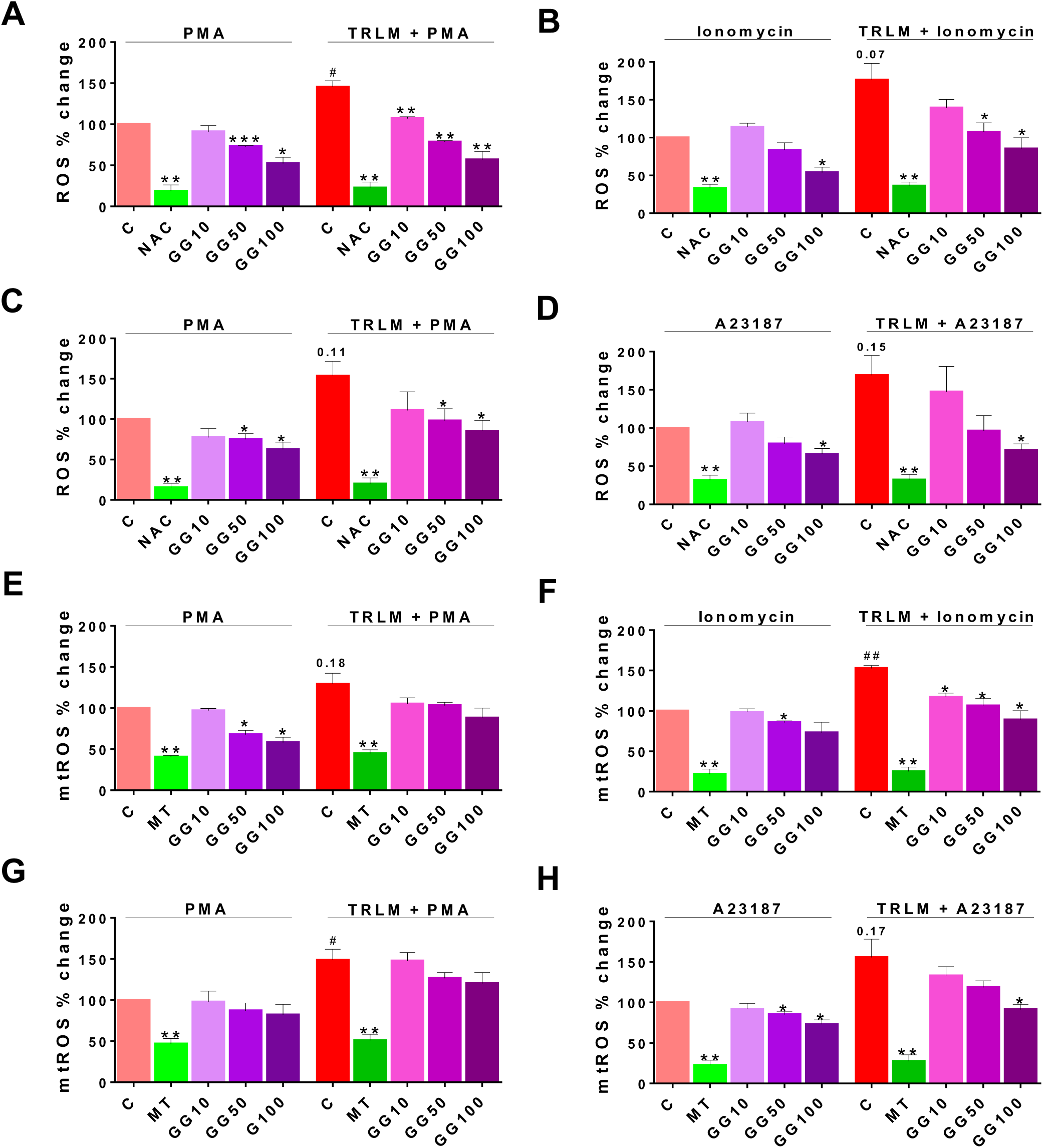
Effect of GG on TRLM primed PMA/ionophores induced NETs formation in human PMNs and murine BMDNs. After pre-incubation with different concentrations GG, cells were treated with TRLM (10 µg/ml) for 30 min and stimulated with sub-maximal concentration of PMA (12.5 nM) and A23187 (1.25 µM) / ionomycin (2 µM) for 30 min. SYTOX Green (100 nM) was used to monitor extracellular DNA release using a plate reader (**A-B**: PMNs; **C-D**: BMDNs). Total MFI in each experimental condition is expressed as Mean ± SEM of 4-6 experiments. A significant increase in NETs formation was induced by TRLM-PMA/ionophores coupling. GG showed a noticeable inhibitory effect on ionophore-induced NETosis. **(E)** NETosis in human PMNs was also monitored using immunofluorescence imaging with DAPI (blue), anti-MPO antibody (green), and anti-H4Cit3 antibody (red). GG showed reduction in nuclear size after induction with ionomycin. Representative fields are shown at 100X with a scale bar of 10 µm. **(E)** Bar diagram representing quantification of percent NETs forming cells as calculated from five transects from three independent experiments. VAS2870 and Diltiazem were used as the positive controls. All the data are represented as Mean ± SEM, n = min 3 per group, and statistical analysis consisted of one-way ANOVA followed by Bonferroni’s test (*p<0.05, **p<0.01, ***p<0.001 vs respective control groups; #p<0.05, ##p<0.01 vs PMA/ionophores treated groups). C, control; V, VAS2870; DTZ, Diltiazem; G10, GG 10 μg/ml; G50, GG 50 μg/ml; G100, GG 100 μg/ml; G300, GG 300 μg/ml; T, TRLM; P, PMA; I, Ionomycin; A, A23187.

Moreover, we further confirmed the anti-NETotic effect of GG *in vitro* by immunofluorescence assay. Primed and induced PMNs exhibited a prominent swollen and diffused cell with thread-like structures (**Fig. 5E**). In agreement with our fluorimetry data, we found comparatively more effect of GG against calcium ionophore treated group. **Fig. 5E & 5F** showed that the pre-treatment of PMNs with 300 μg/ml GG prevented the diffused and web-like state of TRLM-ionomycin treated cells as evident by the reduction of percent NETs forming cells, MPO, and H4Cit3 expression. Incubation of GG with the TRLM-PMA group did not result in much elimination of characteristic DNA fibers extrusion, except with shrinkage of nuclear diameter. Results obtained thus indicate that GG might contribute to the regulation of neutrophil NETs formation via modulating the ionophore mediated signaling pathways involved in NETosis.

### GG suppress ROS and mtROS production

As GG treatment showed a reduction in NETs formation in the primed and activated neutrophils from both mice and human, we therefore examined its anti-oxidant activity in an attempt to decipher its mechanism. When treated with PMA or ionophores before priming with TRLM, a steep rise in superoxide radical production of more than 50% were observed in both the PMNs and BMDNs; ROS (48% for PMA, 75% for ionomycin in human and 55% for PMA, 70% for A23187 in mice) and mtROS (30% for PMA, 51% for ionomycin in human and 50% for PMA, 60% for A23187 in mice) (**Fig. 6A-H).**

Effect of GG on ROS and mtROS production exposed to TRLM-PMA/ ionophores in the presence or absence of GG was measured using flow cytometry. A marked decrease in ROS production in PMNs, from 20% (10 μg/ml), 37% (50 μg/ml) to 50% (100 μg/ml, p<0.05) was observed when stimulated with TRLM-ionomycin whereas TRLM-PMA led to a decrease of 22% at 10 μg/ml, 45% at 50 μg/ml, and 60% at 100 μg/ml, p<0.05 (**Fig. 6A & 6B**). In BMDNs also GG had similar effects in limiting the formation of superoxide anions elicited by TRLM-A23187 (13% reduction at 10 μg/ml, 43% at 50 μg/ml, and 58% at 100 μg/ml, p<0.05) or TRLM-PMA (maximum decrease of 45% at 100 μg/ml, p<0.05) (**Fig. 6C & 6D).** The median inhibitory concentration (IC50) of GG on ROS in BMDNs were 52.3 µg/ml with PMA and 41.8 µg/ml with A23187 (**Fig. S3D & S3E**).

Similarly, GG concentrations were selected for mtROS assay based on the calculation of its IC50 in mice BMDNs; 61.8 μg/ml with PMA and 56.3 μg/ml with A23187 (**Fig S3F & S3G**). The ability of GG, 100 μg/ml to inhibit mtROS production in PMNs revealed a 40% and 30% reduction with TRLM-ionomycin and TRLM-PMA respectively (**Fig. 6E & 6F**). A similar percent reduction was also seen in murine cells with TRLM-A23187 (40%), TRLM-PMA (20%) at 100 μg/ml of GG (**Fig. 6G & 6H).** Notably, GG was comparatively more efficient in reducing calcium ionophores mediated ROS and mtROS production in neutrophils.

One of the hallmark features of COVID-19 is the neutrophilia and cytokine storm which occurs more readily in the lungs as neutrophil numbers are high in pulmonary vasculature than systemic blood vessels (41). We evaluated the degree of inflammation in response to TLR7/8 priming and activation by PMA or A23187 in murine BMDNs, by monitoring the levels of IL-6 and TNFα and their modulation by GG using commercial ELISA kits (**Fig. 4SA & 4SB**). An increase in the production of IL-6 and TNFα were observed when cells were primed with TRLM before treatment with the inducers; IL-6 (52% in PMA and 78% in A23187) and TNFα (96% in PMA and 45% in A23187. GG (100 μg/ml) exerted a profound inhibitory effect on the release of IL-6 and TNFα from neutrophils when treated with either TRLM-PMA or TRLM-A23187. In particular, IL-6 and TNFα levels were reduced to 33% and 37% (p<0.05), respectively when exposed to TRLM-A23187, whereas TRLM-PMA treatment resulted in a decrease of 15-20% in both the cytokines.

Because phagocytosis is one of the important functions of neutrophils to evade any foreign pathogen, we next examined the effect of GG, if any on this property of the neutrophils. Human PMNs pretreated with GG for 60 min followed by PE-labelled latex beads (1 µm) exhibited a modest reduction (14% (p<0.05)) in the phagocytosis only at the highest concentration (300 µg/ml, **Fig. S4C**), while 1-100 µg/ml of GG did not cause any noticeable change. Further, since GG treatment showed mild inhibitory effect on phagocytosis we subsequently assessed its effect on the bactericidal activity. Incubation of overnight grown kanamycin-resistant *E. coli* with neutrophils resulted in the reduction of colonies from 7.9 x 10^8^ to 2.4 x 10^8^ CFU/ml (70% reduction, p<0.05, **Fig. S4D & S4E**). Pre-treatment of human peripheral neutrophils with 300 µg/ml GG did not impart any significant effect on the killing activities of phagocytes; approximately 50% reduction in *E. coli* growth was observed when bacteria were incubated with GG treated PMNs. However, a notable reduction in bacterial growth of 17% (p<0.05) was observed at 300 µg/ml GG, suggesting a direct anti-microbial effect of GG on microbial growth (**Fig. S4D & S4E**).

### Prophylactic use of GG ameliorates COVID-19 pathology in hamster

In summary, we report that the prophylactic use of GG in SARS-CoV-2 infected hamsters helps in mitigating the COVID-19 pathology. The effect of GG against COVID-19 could be through its potent immunomodulatory potential, inhibitory effects on NETosis and suppression of ROS generation as illustrated in the summary figure (**Fig. 7**).

**Figure 7:**
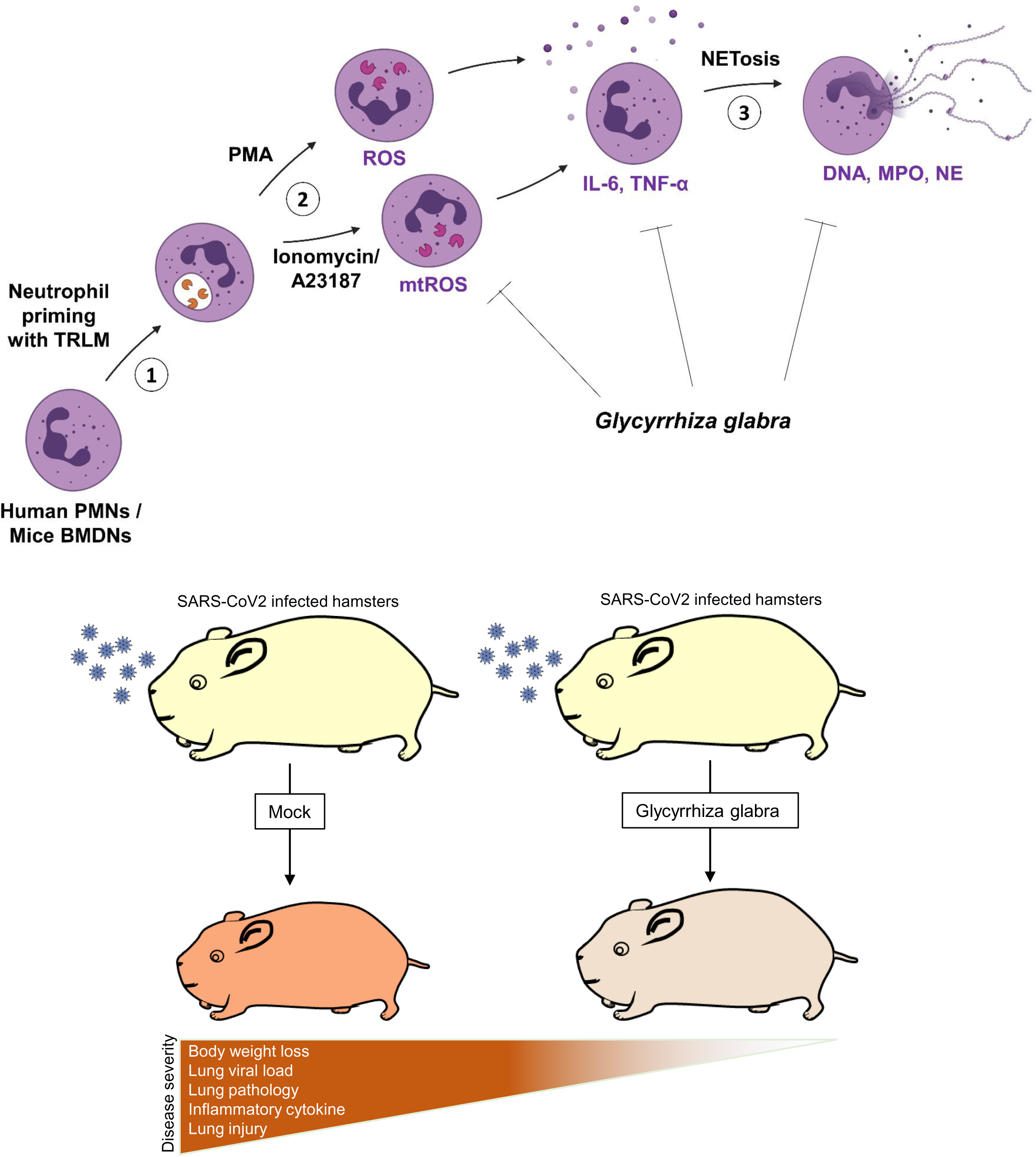
Effect of GG on phagocytosis and pro-inflammatory cytokines release by human PMNs and murine BMDNs. **(A)** Human PMNs were incubated with different concentrations of GG before adding PE-labelled latex beads for phagocytic assay. Fluorescent signal was quenched using trypan blue (0.4%) before acquiring in FACS cell analyzer. GG showed a marginal percent reduction in phagocytic activity. **p<0.01 vs control. **(B-C)** Effect of GG on bactericidal activity of human PMNs. Cells (6.0 x 10^6^) were pre-treated with 300 µg/ml of GG before incubating with kanamycin-resistant *E. coli* (3.0 x 10^8^ CFU/ml). The direct effect of GG on *E. coli* growth was also monitored by incubating bacteria with 300 ug/ml GG for 30 min. **p<0.01 vs control groups; ##p<0.01 vs GG treated groups. **(D-E)** Mice BMDNs were pre-incubation with 100 µg/ml GG and then treated with TRLM (10 µg/ml) for 30 min and stimulated with sub-maximal concentration of PMA (12.5 nM) and A23187 (1.25 µM) for 30 min. Cell supernatant was collected and IL-6 and TNFα concentrations were determined. Significant increases in IL-6 and TNFα levels were induced by TRLM-PMA/A23187 coupling. GG showed a marked inhibitory effect on A23187-induced cytokines secretion. *p<0.05, **p<0.01, vs respective control groups; #p<0.05, ##p<0.01 vs PMA/A23187 treated groups. All the data are represented as Mean ± SEM, n = min 3 per group, and statistical analysis consisted of one-way ANOVA followed by Bonferroni’s test. C, control; GG100, 100 μg/ml; GG300, 300 μg/ml; T, TRLM; P, PMA; A, A23187.

**Figure 8:** Schematic diagram depicting the protective efficacy of prophylactic treatment of GG on SARS-CoV-2 infected hamsters and a putative mechanism of protection.

## Discussion

Recent emergence of SARS-CoV-2 has caused unprecedented mortality which led to development and emergency usage of COVID-19 vaccines (https://covid19.who.int/). Though active vaccination against SARS-CoV-2 has been a largely successful strategy, emerging variants such as Delta (B.1.617.2) and Omicron variant (B.1.1.516) of SARS-CoV-2 have largely been able to escape the immune response elicited by the vaccination (42, 43). This is because of the mutations in the spike/ RBD regions of the variants which were used as an immunogen for several vaccine candidates. Therapeutic use of anti-viral drugs such as Remdesivir (a Rdrp prodrug inhibitor) or Favipiravir (an inhibitor of Rdrp) has been shown to decrease viral load and improve COVID-19 complications (16, 17). However, novel therapeutics development and clinical trials often ends up taking many years which demands repurposing of the existing compounds. Another exciting alterative therapy is provided by herbal medicines which are often used as a food supplement in many places in Asia and Europe (19,20,23). Recent publications on COVID-19 have shown that herbal extracts which exerts immunomodulatory activity maybe beneficial against COVID-19. A recent report published in PNAS 2021 used 3000 Chinese herbal extracts from traditional medicine for the high throughput screening against SARS-CoV-2 in-vitro (23). The study identified a number of potential herbal extracts that showed remarkable ability to inhibit SARS-CoV-2 entry or replication.

In the current study GG anti-viral and immunomodulatory efficacy against SARS-CoV-2 was investigated by using small animal model of SARS-CoV-2 infection. GG is a Indian traditional herb which is part of ancient Ayurveda medicine (44). GG has been shown to contain various active pharmaceutical ingredients such as Glycyrrhizin, liquiritin, isoliguiritin, glyasperin A and glycyrrhizic acid has been shown to have some degree of inhibitory activity against SARS-CoV-2 as shown by computational studies or anti-viral screening *in vitro* (30–33,45). Moreover, a multicomponent study under phase 2 and 3 clinical trials, involving the deglycyrrhizinated form of GG has shown positive results in boosting the host immunity against COVID-19 (46) ClinicalTrials.gov Identifier: NCT04553705). However, till date no studies have been conducted to investigate the protective response of GG against SARS-CoV-2 infection by using animal model.

Corroborating the previously published computational results we found that GG when given as prophylactic regime to hamsters results in substantially reduced gross pathology with significant protection in body weight loss. Moreover, GG group also showed reduction in the lung viral load as compared to the infected group however as compared to the remdesivir group the inhibition by GG was lesser by 10-15%. Protection in the gross clinical parameters of GG administered animals prompted us to look at the pathophysiology of lung as COVID-19 clinical cases are characterized by lung injury and pneumonitis. Remarkably, the histological parameters of GG lungs showed alleviation of lung pathology with overall reduced lung injury score and lowered expression of lung injury markers. These results suggested as that prophylactic treatment of GG is beneficial against COVID-19 in hamster which involves resolving of the pulmonary pathology.

Previously published literature on GG or its components have shown robust immunomodulatory effects of GG. For example, licorice which is a constituent of GG was able to suppress LPS-induced NLRP3 inflammasomes and NF-kB activation (47). This would mean that GG could lower the inflammatory response induced by LPS. When the mechanism of this inhibition was investigated, it was found that licorice inhibits COX-2 expression and in turn reduces the prostaglandin (PGE2) levels (28, 47). Another study have documented that glycyrrhizin, another component of GG, reduces ROS generation by neutrophils thereby suppressing tissue inflammation (27, 48). These studies prompted us also to investigate the immunomodulatory potential of GG. When spenocytes mRNA expression was studied for cytokines, we found that GG remarkably inhibited pro-inflammatory cytokines in a way which was comparable to remdesivir, a prototype of anti-viral drug. Higher levels of cytokines such as TNFα, IL17 in the early stages of infection have been linked with worsening of the disease and tissue injury. Remarkably, GG was able to suppress the expression of all these 3 cytokines at the peak of infection in hamster (ie 4dpi). In addition, GG also suppressed the expression of T-bet which is a key transcription factor for Th1 response. This was an interesting finding, corroborating the immunomodulatory potential of GG previously shown in the context of other pathogenic infection, as it proved that the protective effect of GG against COVID-19 involves in parts both anti-viral activity as well as immunomodulatory potential resulting in anti-inflammatory response.

CD4+ T helper cells are crucial components of adoptive immunity which coordinates in providing immunity against pathogenic infection. On the other hand, it has been also shown that dysregulated effector and regulatory T helper cell response leads to tissue injury and worsening of the disease which is also seen in the case of COVID-19 (37, 49). To further validate the anti-inflammatory potential of GG and to evaluate GG effect on suppression of effector T cells, we performed T cell differentiation assay in-vitro in presence or absence of GG. Our results show that GG could inhibit the differentiation of Th1, Th2 and Th17 differentiation in a dose dependent manner much like the dexamethasone, a known immunosuppressant (29). Interestingly, the lowest ID50 for in-vitro assay was found to be for Th1 differentiation suggesting that GG at lower doses inhibits Th1 differentiation dramatically. Since GG was also found to significantly inhibit thrombosis marker PAI-1, and during COVID-19 induction of thrombosis is strongly correlated with the neutrophils induced NETosis, we investigated further if GG could modulate the neutrophil response and affect ROS generation. While neutrophils are an important innate immune cells in the context of viral immunity, generation of ROS has been linked with cellular injury leading to pulmonary damage and extra pulmonary organ damage (10). NETs are made up of the dsDNA fibers extruded from neutrophils, containing citrullinated histones and granular enzymes, such as myeloperoxidase (MPO), neutrophil elastase (NE), cathepsin G, α-defensins, bactericidal permeability-increasing factor (BPI), and pentraxin 3. Agonists of endosomal TLR 7/8 which binds to viral single-stranded RNA as in SARS-Co-2, influenza virus have been shown to induce neutrophil activation, however little is known about a putative link between TLR7/8 signaling and the release of NETs in neutrophils. In the present study TRLM upregulated PMA, and calcium ionophores induced neutrophil functions, indicative of its priming effect. Though the effect of TRLM on oxidative stress in neutrophils has not been elucidated before, similar agonists such as resiquimod (R848) and its water-soluble derivative CL097, have been found to prime the neutrophils and induce cytokines and ROS production (50–52). The molecular basis of such enhancement of neutrophil-derived functions is still lacking, however priming of TLR7/8 receptors enhances the survival of neutrophils and also augment the neutrophil functions (53–55).

Involvement of TLRs for the release of NETs has already been described earlier; Awasthi *et al*. (2016) had reported that blocking TLR-2 and -6 with specific antibodies significantly reduced oxLDL induced NETs formation (56). Our in-vitro data highlighted that NETs released in TRLM primed PMNs following treatment with PMA and calcium ionophore, were reduced by GG pretreatment in a concentration-dependent manner. The amount of DNA release in neutrophils stimulated with calcium ionophores showed a more robust reduction as also evident by immunolabeling of MPO and citrullinated histones in the GG pretreated cells. There are no reports on the effect of GG on NETs, however other medicinal herbs such as Danshen, the dried root of *Salvia miltiorrhiza*, and the compounds like salvianolic acid B and 15,16-dihydrotanshinone have been found to be effective in reducing NETs by inhibiting MPO and NOX (57). Extracts from *Eugenia aurata* and *E. punicifolia* also inhibited inflammatory response by reducing neutrophil adhesion, degranulation, and NETs release (58).

Classically PMA mediated NOX-dependent ROS generation is mediated by the activation of protein kinase C (59, 60) while NOX-independent pathway is mediated by calcium-activated small conductance potassium channel (SK3) and/or non-selective mitochondrial permeability transition pore (mPTP) in inducing mtROS production via intracellular Ca^2+^ flux (61, 62). Importantly, numerous reports have described the crosstalk of mtROS production and NOX activation representing a feed-forward vicious cycle of ROS production in oxidative stress (63). Douda *et al*. (2015) failed to observe cytoplasmic ROS generation via NOX in the presence of A23187 (61). While Vorobjeva *et al*. (2020) by using mitochondrial-targeted antioxidant and inhibitors of NOX showed involvement of both NOX-derived ROS and mtROS in calcium ionophore induced NETosis in human PMNs (62). We also observed a large amount of mtROS production by ionomycin during NOX-independent NETosis. We reported a significant reduction in the oxidative stress in GG pretreated cells as demonstrated by its ability to suppress both ROS and mtROS. However, the anti-NETotic effect of GG was comparatively more pronounced in case of ionophores than PMA. This indicates a putative role of GG as a mitochondrial-targeted anti-oxidant in preventing the mtROS - NOX feed-forward cycle and the ensuing oxidative insult and NETosis. Recently Fortner *et al*. (2020) have also found targeting mitochondrial oxidative stress with mitochondrial-targeted anti-oxidant MitoQ, reduced neutrophil mtROS and NETs formation in lupus-prone mice (64). Although our result point out mtROS as the prominent player, however, we cannot rule out the possibility of GG in reducing NETosis through other signaling pathways, particularly NOX2-ROS axis, which require further work to decode the detailed signaling pathways.

Instillation of Ayurvedic herb, Anu oil exhibited reduced SARS-CoV-2 load and the expression of pro-inflammatory cytokine genes (Th1 and Th17) in the lungs (38). Recent reports have shown upregulation in IL-6 from human neutrophils using TLR8 ligand, resiquimod, and IFNα (65). NF-κB activation and the associated signaling seems to be the major pathways activated downstream of TRL7/8 activation leading to the release of pro-inflammatory cytokines. In one study using monocyte, Singh *et al.* (2015) had revealed enhanced ROS production and increased transcription, processing, and secretion of IL-1β upon activation of TLR-2 and -4 via cross-talk of p67phox-NOX2 with IRAK-ERK pathway (66). ROS-dependent pathway for the activation of inflammatory response via NOD-like receptor protein 3 (NLRP3) inflammasome has also been established (67, 68). Further, multiple reports have suggested the involvement of NFκB in the regulation of the NLRP3 inflammasome (69). PAD enzymes are calcium-dependent enzymes and PAD4 plays a prime role in ionomycin/A23187-dependent NETosis. Interestingly PAD4 promotes NETs via regulating the assembly of NLRP3 inflammasome (70). Our results corroborate with this notion as A23187 showed significantly higher up-regulation in IL-6 secretion as compared to PMA. Owing to the significant inhibition of ROS production, the anti-inflammatory effects of GG are apparent. The protective effect of GG on TRLM-primed inflammation showed a higher efficacy against calcium ionophore, further pointing out the critical role of NOX-independent pathways in GG-mediated regulation of different neutrophil functions. Although GG plays a role in the modulation of cytokines expression and its influence on NETs has been explored, the mechanism involved remains to be elucidated.

Besides NETosis to control the pathogen infection, these granulocytes have a plethora of other defensive mechanisms. Being the most abundant leukocyte in blood, neutrophils are the prime phagocytes of the innate immune system (71). Modulation of innate immunity by medicinal plants or by their constituents has seen a renaissance in recent times. Our data did not reveal any major change in the phagocytosis by neutrophils when incubated with GG, however other herbal plants from the same family reported an increase in the phagocytic activity of neutrophils, such as *Caesalpinia bonducella* and *Vigna mungo* (72, 73). Moreover, the intracellular bactericidal capacity of neutrophils was not affected by GG; a mild reduction probably suggests an inhibitory effect due to ROS/RNS production within the phagolysosomes during oxidative burst (74, 75).

In summary, here we provide the first direct evidence based on in-vivo and in-vitro data from hamster, mice and human studies to show potent protective efficacy of GG against COVID-19 pathologies and show that the mechanism of this protection could be mediated by strong anti-inflammatory response, inhibition of NETosis and suppression of ROS generation.

## Methods

GG extract used in the study was prepared as per pharmacopeial standards and was provided by National Medicinal Plant Board for the study.

### Animal Ethics and biosafety statement

6-9 weeks golden Syrian hamsters were procured from CDRI and quarantined for one week at small animal facility (SAF), THST before starting the experiment. Prophylactic GG group started received twice daily oral doses of GG 130 mg/ kg (0.5% CMC preparation) 5 days prior to challenge and continued till end point. Remdesivir control group received 15mpk sc injections of remdesivir on 1 day before and 1 day after the challenge (24, 38). On the day of challenge the animals were shifted to ABSL3. Intranasal infection of live SARS-CoV2 (SARS-Related Coronavirus 2, Isolate USA-WA1/2020)10^5^PFU/ 100μl or with DMEM mock control was established with the help of catheter under mild anesthetized by using ketamine (150mg/kg) and xylazine (10mg/kg) intraperitoneal injection inside ABSL3 facility. All the experimental protocols involving the handling of virus culture and animal infection were approved by RCGM, institutional biosafety and IAEC (IAEC/THSTI/105) animal ethics committee.

### Preparation of GG extract

GG extract was prepared by dissolving 10 mg of dry roots powder of GG in 1 ml of water overnight in shaker incubator at 37°C. Next day, the suspension was centrifuged at 10000 x g for 30 min followed by filtration of supernatant using 0.45 filter. The filtrate was considered as 100% aqueous extract and diluted in water according to the experimental requirements.

### Virus culture and titration

SARS-Related Coronavirus 2, Isolate USA-WA1/2020 virus was grown and titrated in Vero E6 cell line cultured in Dulbecco’s Modified Eagle Medium (DMEM) complete media containing 4.5 g/L D-glucose, 100,000 U/L Penicillin-Streptomycin, 100 mg/L sodium pyruvate, 25mM HEPES and 2% FBS. The stocks of virus were plaque purified at THSTI IDRF facility inside ABSL3 following institutional biosafety guidelines.

### Gross clinical parameters of SARS-CoV2 infection

Post challenge, the body weight of the animals was recorded daily till 4 day post infection (4 dpi). On 4 dpi all animals were euthanized at ABSL3 and necropsy was performed to collect organs. Lungs and spleen of the animals were excised and imaged for gross morphological changes (24). Left lower lobe of the lung was fixed in 10% formalin and used for histological analysis (76). The remaining part of the lung left lobe was homogenized in 2ml Trizol solution for viral load estimation. Spleen was homogenized in 2ml of Trizol solution. The trizol samples were stored immediately at -80 °C till further use. Serum samples isolated from blood was immediately stored at -80 °C till further use.

### Viral load

Isolated lung was homogenized in 2ml Trizol reagent (Invitrogen) and RNA was isolated by Trizol-Choloform method. Yield of RNA yield was quantitated by nano-drop and 1 µg of RNA was use to reverse-transcribed to cDNA using the iScript cDNA synthesis kit (BIORAD; #1708891) (Roche). 1:5 diluted cDNAs was used for qPCR by using KAPA SYBR® FAST qPCR Master Mix (5X) Universal Kit (KK4600) on Fast 7500 Dx real-time PCR system (Applied Biosystems) and the results were analyzed with SDS2.1 software (24, 38). Briefly, 200 ng of RNA was used as a template for reverse transcription-polymerase chain reaction (RT-PCR). The CDC-approved commercial kit was used for of SARS-CoV-2 N gene: 5′-GACCCCAAAATCAGCGAAAT-3′ (Forward), 5′-TCTGGTTACTGCCAGTTGAATCTG-3′ (Reverse). Hypoxanthine-guanine phosphoribosyltransferase (HGPRT) gene was used as an endogenous control for normalization through quantitative RT-PCR. The relative expression of each gene was expressed as fold change and was calculated by subtracting the cycling threshold (Ct) value of hypoxantine-guanine phosphoribosyltransferase (HGPRT-endogenous control gene) from the Ct value of target gene (ΔCT). Fold change was then calculated according to formula POWER(2,-ΔCT)*10,000 (77, 78).

### qPCR from splenocytes

RNA isolated from spleen samples were converted into cDNA as described above. Thereafter, the relative expression of each gene was expressed as fold change and was calculated by subtracting the cycling threshold (Ct) value of hypoxantine-guanine phosphoribosyltransferase (HGPRT-endogenous control gene) from the Ct value of target gene (ΔCT). Fold change was then calculated according to formula POWER(2,-ΔCT)*10,000. The list of the primers is provided as follows.

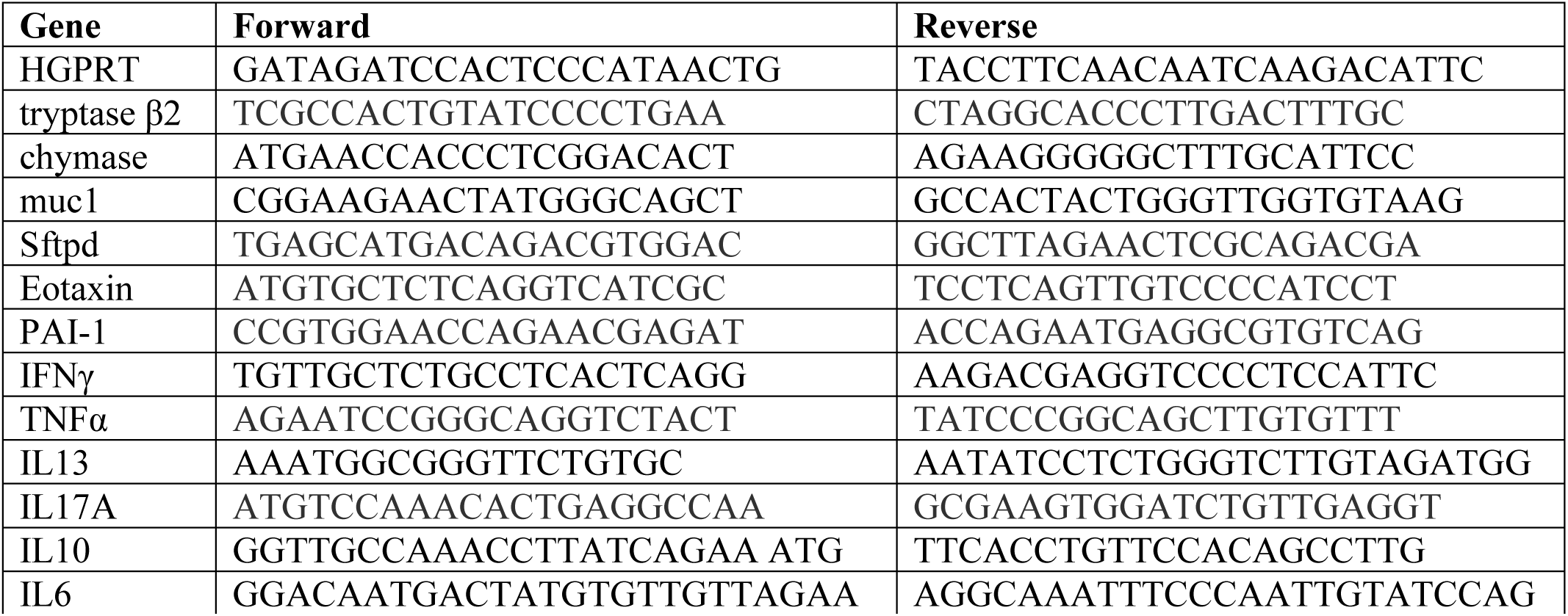

### Histology

Fixed lung samples were embedded in paraffin blocks and sectioned into 3 µm sections mounted on silane-coated glass slides. The sectioned slides were then used for staining with hematoxylin and eosin dye as previously described (76). Each stained sample were then analyzed and captured at 10X magnification. Assessment for the histological score was carried out through blind scoring for each sample by a professional histologist on a scale of 0-5 (where 0 indicated absence of histological feature while 5 indicated highest score). Disease index score was calculated by addition of all the individual histological scores.

#### In vitro differentiation of T cells

Single cell suspension was prepared from spleen and lymph nodes of 6–8 weeks old C57BL/6 mice. The cells were activated using soluble anti-CD3 (2ug/ml) and differentiated into Th1 conditions by adding recombinant mouse IL-12 (15ng/ml) cytokine or Th2 conditions by adding recombinant mouse IL-4 (15ng/ml) cytokine or Th17 conditions by adding TGF-beta (2ng/ml) plus IL-6 cytokine (25ng/ml) (78, 79). *Glycyrrhiza gabra* was added in concentrations ranging from 10ug/ml to 1000ug/ml at the start of culture. Cells were harvested after 72 hours of culture. Intracellular cytokine staining was performed to check expression of IFNγ, IL-4 and IL-17 cytokine for Th1, Th2 and Th17 cells respectively.

#### Intracellular cytokine staining

Cells were stimulated for 4 h with PMA (phorbol 12- myristate13-aceate; 50 ng/ml) and ionomycin (1.0 ug/ml) and a protein-transport inhibitor containing monensin before detection by staining with antibodies. Surface markers were stained for 15–20 min in room temperature in PBS with 1% FBS, then were fixed in Cytofix and permeabilized with Perm/Wash Buffer using Fixation Permeabilization solution kit and stained anti-IL-17A; anti-IFNγ, anti-IL-4 diluted in Perm/Wash buffer. All antibodies were used in 1:500 dilution. The cells were then taken for flow cytometry using BD FACSCantoII and data was analysed with FlowJo software.

### Isolation of murine BMDNs and human peripheral neutrophils

Murine bone marrow-derived neutrophils were isolated from femur and tibia bones of C57BL/6 wild-type male mice (20–25 g, 12–16 weeks) using the method described previously (71). Long bones were flushed with HBSS + 0.1% BSA through a sterile tube followed by centrifugation at 400 x g for 10 min at 10°C. Pellet were resuspended in 45% Percoll and gently layered over Percoll density gradient (81%-62%-55%). After centrifugation at 1700 x g for 30 min with acceleration 5 m/s^2^ and deceleration 4 m/s^2^, band between 81% and 62% were harvested and assessed for their viability by Trypan blue and purity by anti-Ly6G and anti-CD11b antibodies. Human neutrophils were isolated from the peripheral blood obtained from a healthy donor, by following Percoll density gradient method as described earlier (80). Briefly, 2 ml buffy coat were incubated with 6% dextran at 37°C for 30 min. The upper phase was collected and centrifuged at 1800 x g for 10 min. Pellet was mixed in HBSS containing 0.1% BSA and layered on top of the Percoll gradient (81%-62%) before centrifuging at 1800 x g for 30 min. Neutrophils containing layer was collected and purity and viability of neutrophils were assessed by CD15 labeling and trypan blue, respectively. All the studies on mice were approved by the institutional animal (THSTI/105) and human (THS1.8.1/100) ethical committees, DBT-THSTI, Faridabad.

### Cell viability assay

1.0×10^6^ per ml murine BMDNs and human PMNs were incubated with 100-1000 µg/ml of GG for up to 240 min in flow tubes to assess the cytotoxicity of the herbal extracts as described earlier (81). For the cell death analysis, neutrophils were stained with cell-impermeant DNA intercalating propidium iodide (PI, 50 μg/ml) and were kept in dark at RT for 15 min. DNA content was analyzed on BD FACS Canto cell analyzer using BD FACS Diva software (BD Biosciences, USA).

### Intracellular ROS and mtROS analysis

Cytosolic and mitochondrial ROS was measured using the fluorescent probes DCFH-DA (10 μM) and MitoSOX (10 μM), respectively in mouse BMDNs and human PMNs. 1.0 x 10^6^ cells/ml were pre-incubated with GG before treatment with different interventions such as TRLM (10 μM), PMA (10-100 nM), A23187 (1-5 μM), ionomycin (1-4 μM), NAC (10 μM), and MitoTEMPO (10 μM). The control experiment contained the concentration of DMSO (0.1%) as in the extract-treated cells. Minimum 10,000 events were acquired for each sample using BD FACS Canto II.

### NETosis assay

Neutrophils (5.0 x 10^5^ cells/ml) from mice or human were added to poly-L-lysine coated wells and incubated with GG for 60 min at 37°C. SYTOX Green (100 nM) was added to each well, and cells were treated with TRLM (10 uM), PMA (10 uM), A23187 (1 uM), ionomycin (1 uM), VAS2870 (10 μM), Diltiazem (10 μM), or vehicle (DMSO 0.1%) and incubated in a CO_2_ incubator at 37°C. The fluorescence was measured at different time periods up to 240 min in a fluorescence plate reader at 37°C (Synergy 2; BioTek). The fluorescence of stimulated cells was expressed in arbitrary units and was determined by subtracting the baseline fluorescence of unstimulated cells from the activated cells. In a parallel experiment, immunofluorescence staining of BMDNs and PMNs was carried out using mouse anti-MPO and rabbit anti-H4Cit3 antibodies. After fixation and blocking, samples were incubated overnight with 1:100 dilution of primary antibodies and were visualized after incubation with the secondary antibodies (1:200, anti-mice AF488 and anti-rabbit AF594) using the confocal microscope (Olympus FV3000) at 100X resolution. DNA was stained with DAPI (0.5 μg/ml).

### Pro-inflammatory cytokines assay

The levels of IL6 and TNFα in the supernatant/cell-free samples after the centrifugation were measured by using the monoclonal antibody-based mouse interleukin ELISA kits. Briefly 100 μl of media from cells (1.0 x 10^6^ cells/ml) treated with TRLM and PMA/A23187, following pre-incubation with GG/vehicle, was used to measure the cytokines referring the manufacturer’s instruction.

### Phagocytosis and bactericidal assay

To assess the phagocytosis, PE-labelled latex beads were incubated with GG pre-treated human PMNs (1.0 x 10^4^) at 1:50 (neutrophil: beads) ratio and were analyzed using FACS (75) after quenching the extracellular fluorescent signal of adherent beads with Trypan blue (0.4%). In a separate experiment, GG pre-exposed human neutrophils were treated with kanamycin-resistant *E. coli* for 30 min at 37°C. After lysis of the cells using sterile water, the diluted lysate was spread on LB agar plates with the antibiotic and kept at 37°C overnight. The bacterial colonies were counted and the bactericidal activity is expressed as a percent of CFU in the presence and absence of neutrophils.

### Statistical analysis

All the experiments have been carried out independently in triplicate. Results are being expressed as mean ± SEM. Multiple group comparisons have been performed using one-way ANOVA followed by the Bonferroni test using GraphPad Prism 8. The differences have been considered as statistically significant when the *p*-value was < 0.05.

## Acknowledgments

MD and AA received financial support for this study from Ayush-DBT (BT/PR40378/TRM/120/486/2020 & NMPB/IFD/GIA/NR/PL/2020-21/53). AA and MD received financial support from THSTI Core to establish the hamster model for SARS-CoV2 infection. We greatly acknowledge the support and critical inputs of Dr. Pramod Kumar Garg in the manuscript. We thank IDRF (THSTI) for the support at ABSL3 facility. Prabhanjan and Jitender for providing technical support. Small animal facility and Immunology Core for providing support in experimentation. Hamsters were procured from CDRI, Lucknow. ILBS for support in histological analysis and assessment. The following reagent was deposited by the Centers for Disease Control and Prevention and obtained through BEI Resources, NIAID, NIH: SARS Related Coronavirus 2, Isolate USA-WA1/2020, NR-52281.

## Author Contributions

Conceived, designed and supervised the study: MD, AA; Performed the experiments: ZAR, PB, SS, UM; ABSL3 experiment: ZAR, SS; FACS: PB, UM; qPCR: ZAR, PB; Viral load: ZAR; Histology: ZAR; ELISA: PB; Fluorescence microscopy: PB; Analyzed the data: ZAR, PB; Contributed reagents/materials/analysis tools: MD, AA; Wrote the manuscript: ZAR, PB, AA, MD.

## Declaration of Interests

The authors declare no conflict of interest.

## Competing Interests

The authors declare no competing interest.

## Supplementary figure legends

**Supplementary Figure 1:**
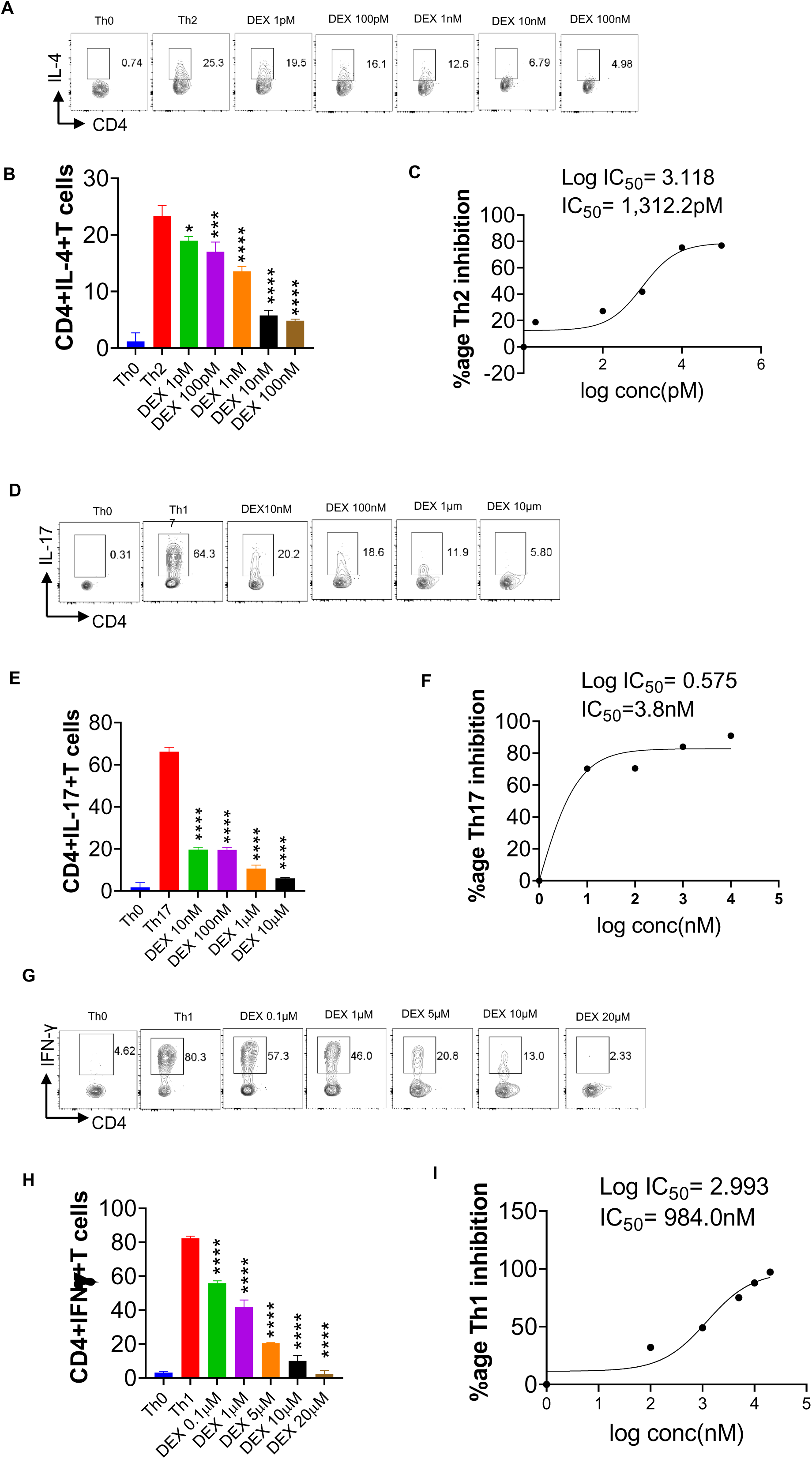
Effect of Dexamethasone on *in vitro* differentiation of Th1, Th2 and Th17 cells. Spleen and Lymph nodes were isolated from 6-8 weeks old C57BL/6 mice and their single cell suspension was prepared. Cells were activated using soluble anti-CD3 antibody and differentiated into helper T (Th) 2 (**A, B**), Th17 cells (**D, E)** and Th1 conditions (**G, H**) using recombinant mouse IL-4; TGF-β + IL-6 and IL-12 cytokines respectively. Dexamethasone was added in doses ranging from 1pM to 20uM at the start of culture. Cells were differentiated for 72 hours and IL-4, Il-17 and IFNγ production was measured by Intracellular cytokine staining. IC50 values were calculated using Graph pad prism software (**C, F, I**). *P<0.05, **P<0.01, ***P<0.001, ****P<0.0001 by one-way ANOVA.

**Supplementary Figure 2:**
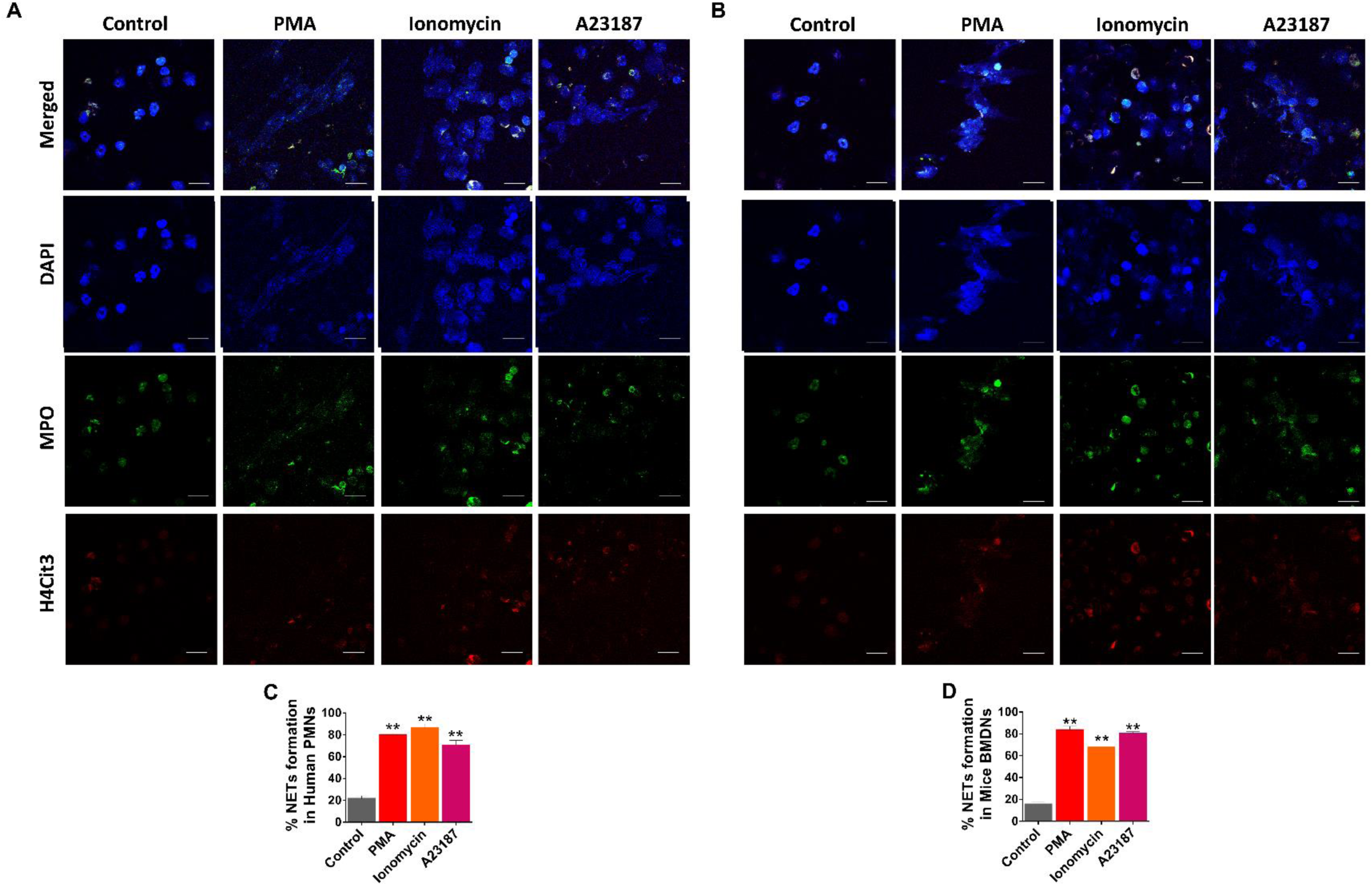
Induction of NETs by PMA and calcium ionophores in human PMNs (A) and mice BMDNs (B). Neutrophils were incubated with PMA (100 nM), A23187 (10 µM), and ionomycin (10 µM), and the release of NETs were followed for up to 240 min and visualized by immunofluorescence imaging. Cells were stained with blue (DNA), green (MPO), and red (H4Cit3). Magnification 100X; Scale bar 10 µm. **(C-D)** Bar graphs represents the quantification of percent NETs forming cells in the presence of PMA, Ionomycin, and A23187. Data presented as percent NETs formation are mean ± SEM of five fields from three independent experiments.

**Supplementary Figure 3:**
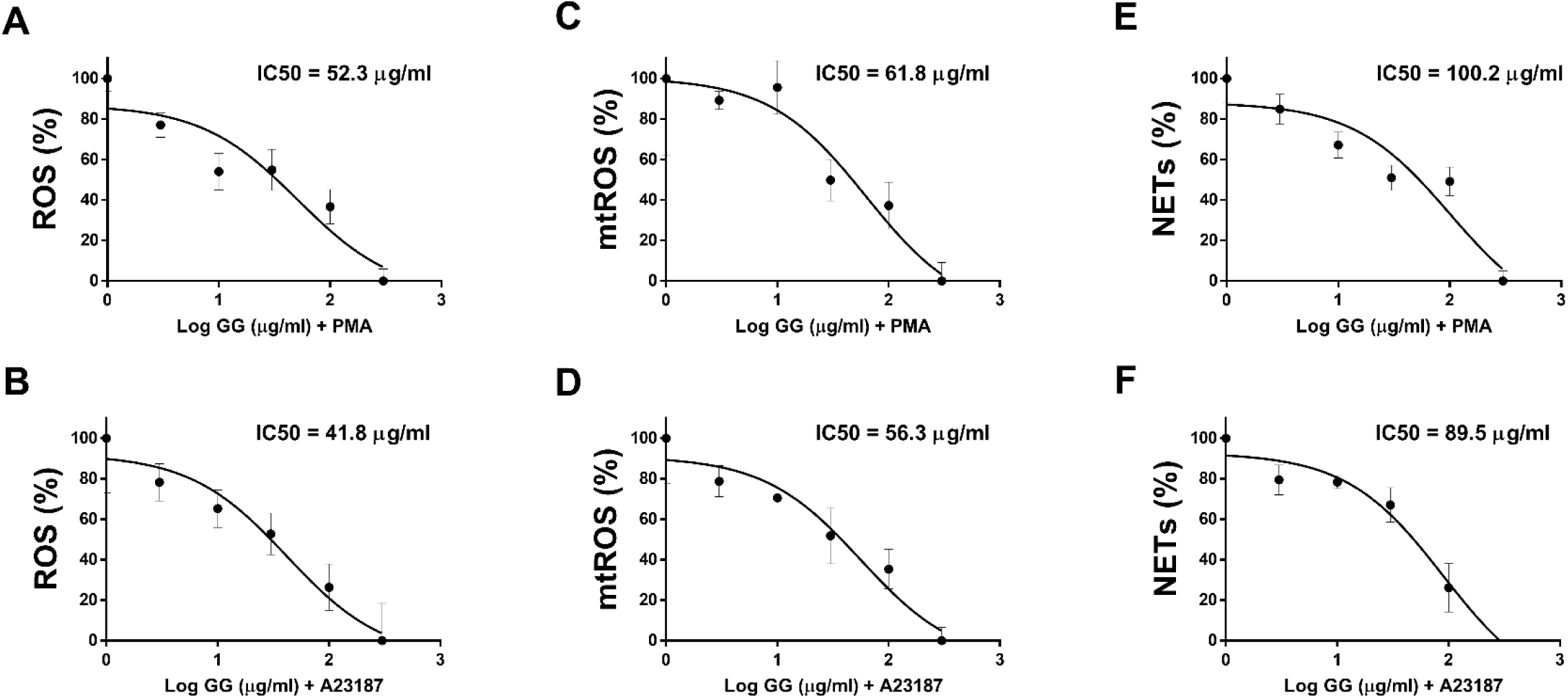
IC50 values of GG on (A-B) ROS, (C-D) mtROS, and (E-F) NETosis in murine BMDNs in the presence of PMA and A23187 as inducers.

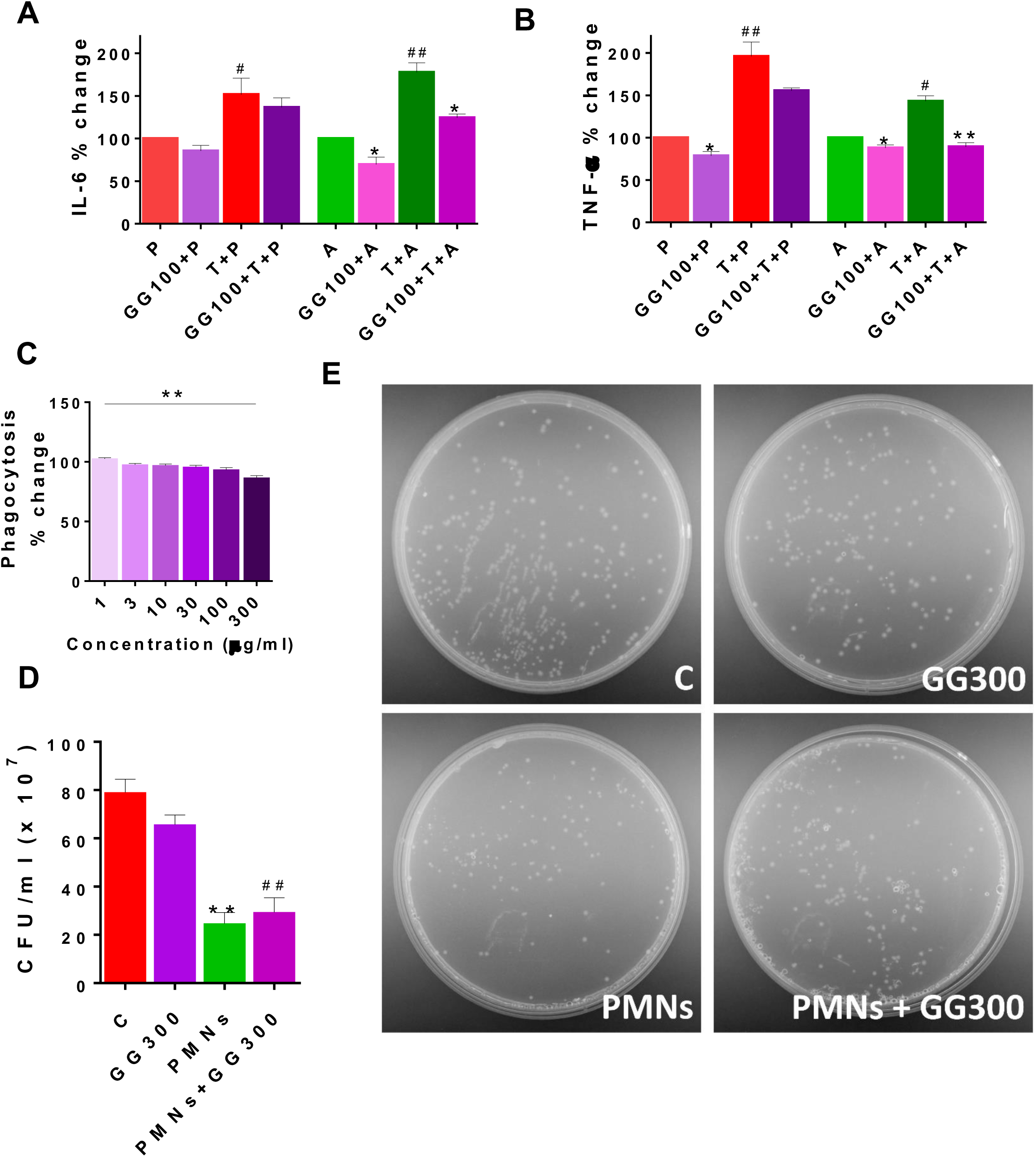

## Notes

### Competing Interest Statement

The authors have declared no competing interest.

## References

1. Chen Y, Li L. SARS-CoV-2: virus dynamics and host response. The Lancet Infectious Diseases (2020) 20:515–516. doi: 10.1016/S1473-3099(20)30235-8

2. Guan W-J, Ni Z-Y, Hu Y, Liang W-H, Ou C-Q, He J-X, Liu L, Shan H, Lei C-L, Hui DSC, et al. Clinical Characteristics of Coronavirus Disease 2019 in China. N Engl J Med (2020) 382:1708–1720. doi: 10.1056/NEJMoa2002032

3. Moore JB, June CH. Cytokine release syndrome in severe COVID-19. Science (2020) 368:473–474. doi: 10.1126/science.abb8925

4. Tan L, Wang Q, Zhang D, Ding J, Huang Q, Tang Y-Q, Wang Q, Miao H. Lymphopenia predicts disease severity of COVID-19: a descriptive and predictive study. Sig Transduct Target Ther (2020) 5:1–3. doi: 10.1038/s41392-020-0148-4

5. Nishiga M, Wang DW, Han Y, Lewis DB, Wu JC. COVID-19 and cardiovascular disease: from basic mechanisms to clinical perspectives. Nature Reviews Cardiology (2020) 17:543–558. doi: 10.1038/s41569-020-0413-9

6. Yang L, Tu L. Implications of gastrointestinal manifestations of COVID-19. The Lancet Gastroenterology & Hepatology (2020) 5:629–630. doi: 10.1016/S2468-1253(20)30132-1

7. Zhong P, Xu J, Yang D, Shen Y, Wang L, Feng Y, Du C, Song Y, Wu C, Hu X, et al. COVID-19-associated gastrointestinal and liver injury: clinical features and potential mechanisms. Signal Transduction and Targeted Therapy (2020) 5:1–8. doi: 10.1038/s41392-020-00373-7

8. Mao L, Jin H, Wang M, Hu Y, Chen S, He Q, Chang J, Hong C, Zhou Y, Wang D, et al. Neurologic Manifestations of Hospitalized Patients With Coronavirus Disease 2019 in Wuhan, China. JAMA Neurol (2020) 77:683–690. doi: 10.1001/jamaneurol.2020.1127

9. Liu J, Liu Y, Xiang P, Pu L, Xiong H, Li C, Zhang M, Tan J, Xu Y, Song R, et al. Neutrophil-to-lymphocyte ratio predicts critical illness patients with 2019 coronavirus disease in the early stage. Journal of Translational Medicine (2020) 18:206. doi: 10.1186/s12967-020-02374-0

10. Arcanjo A, Logullo J, Menezes CCB, de Souza Carvalho Giangiarulo TC, dos Reis MC, de Castro GMM, da Silva Fontes Y, Todeschini AR, Freire-de-Lima L, Decoté-Ricardo D, et al. The emerging role of neutrophil extracellular traps in severe acute respiratory syndrome coronavirus 2 (COVID-19). Sci Rep (2020) 10:19630. doi: 10.1038/s41598-020-76781-0

11. Zhu Y, Chen X, Liu X. NETosis and Neutrophil Extracellular Traps in COVID-19: Immunothrombosis and Beyond. Frontiers in Immunology (2022) 13: https://www.frontiersin.org/article/10.3389/fimmu.2022.838011 [Accessed May 11, 2022]

12. Veras FP, Pontelli MC, Silva CM, Toller-Kawahisa JE, de Lima M, Nascimento DC, Schneider AH, Caetité D, Tavares LA, Paiva IM, et al. SARS-CoV-2–triggered neutrophil extracellular traps mediate COVID-19 pathologySARS-CoV-2 directly triggers ACE- dependent NETs. Journal of Experimental Medicine (2020) 217:e20201129. doi: 10.1084/jem.20201129

13. Kimball AS, Obi AT, Diaz JA, Henke PK. The Emerging Role of NETs in Venous Thrombosis and Immunothrombosis. Frontiers in Immunology (2016) 7: https://www.frontiersin.org/article/10.3389/fimmu.2016.00236 [Accessed May 11, 2022]

14. Dong Y, Dai T, Wei Y, Zhang L, Zheng M, Zhou F. A systematic review of SARS-CoV-2 vaccine candidates. Signal Transduction and Targeted Therapy (2020) 5:1–14. doi: 10.1038/s41392-020-00352-y

15. Li S, Chen C, Zhang H, Guo H, Wang H, Wang L, Zhang X, Hua S, Yu J, Xiao P, et al. Identification of natural compounds with antiviral activities against SARS-associated coronavirus. Antiviral Research (2005) 67:18–23. doi: 10.1016/j.antiviral.2005.02.007

16. Beigel JH, Tomashek KM, Dodd LE, Mehta AK, Zingman BS, Kalil AC, Hohmann E, Chu HY, Luetkemeyer A, Kline S, et al. Remdesivir for the Treatment of Covid-19 — Final Report. New England Journal of Medicine (2020) 383:1813–1826. doi: 10.1056/NEJMoa2007764

17. Kaptein SJF, Jacobs S, Langendries L, Seldeslachts L, Horst S ter, Liesenborghs L, Hens B, Vergote V, Heylen E, Barthelemy K, et al. Favipiravir at high doses has potent antiviral activity in SARS-CoV-2−infected hamsters, whereas hydroxychloroquine lacks activity. PNAS (2020) 117:26955–26965. doi: 10.1073/pnas.2014441117

18. Jan J-T, Cheng T-JR, Juang Y-P, Ma H-H, Wu Y-T, Yang W-B, Cheng C-W, Chen X, Chou T-H, Shie J-J, et al. Identification of existing pharmaceuticals and herbal medicines as inhibitors of SARS-CoV-2 infection. PNAS (2021) 118: doi: 10.1073/pnas.2021579118

19. Li S, Cheng C-S, Zhang C, Tang G-Y, Tan H-Y, Chen H-Y, Wang N, Lai AY-K, Feng Y. Edible and Herbal Plants for the Prevention and Management of COVID-19. Front Pharmacol (2021) 12: doi: 10.3389/fphar.2021.656103

20. Matveeva T, Khafizova G, Sokornova S. In Search of Herbal Anti-SARS-Cov2 Compounds. Front Plant Sci (2020) 11: doi: 10.3389/fpls.2020.589998

21. Panyod S, Ho C-T, Sheen L-Y. Dietary therapy and herbal medicine for COVID-19 prevention: A review and perspective. Journal of Traditional and Complementary Medicine (2020) 10:420–427. doi: 10.1016/j.jtcme.2020.05.004

22. Zhang D, Wu K, Zhang X, Deng S, Peng B. In silico screening of Chinese herbal medicines with the potential to directly inhibit 2019 novel coronavirus. Journal of Integrative Medicine (2020) 18:152–158. doi: 10.1016/j.joim.2020.02.005

23. Jan J-T, Cheng T-JR, Juang Y-P, Ma H-H, Wu Y-T, Yang W-B, Cheng C-W, Chen X, Chou T-H, Shie J-J, et al. Identification of existing pharmaceuticals and herbal medicines as inhibitors of SARS-CoV-2 infection. PNAS (2021) 118: doi: 10.1073/pnas.2021579118

24. Rizvi ZA, Dalal R, Sadhu S, Binayke A, Dandotiya J, Kumar Y, Shrivastava T, Gupta SK, Aggarwal S, Tripathy MR, et al. Golden Syrian hamster as a model to study cardiovascular complications associated with SARS-CoV-2 infection. eLife (2022) 11:e73522. doi: 10.7554/eLife.73522

25. Chan JF-W, Zhang AJ, Yuan S, Poon VK-M, Chan CC-S, Lee AC-Y, Chan W-M, Fan Z, Tsoi H-W, Wen L, et al. Simulation of the Clinical and Pathological Manifestations of Coronavirus Disease 2019 (COVID-19) in a Golden Syrian Hamster Model: Implications for Disease Pathogenesis and Transmissibility. Clinical Infectious Diseases (2020) 71:2428–2446. doi: 10.1093/cid/ciaa325

26. Sia SF, Yan L-M, Chin AWH, Fung K, Choy K-T, Wong AYL, Kaewpreedee P, Perera RAPM, Poon LLM, Nicholls JM, et al. Pathogenesis and transmission of SARS-CoV-2 in golden hamsters. Nature (2020) 583:834–838. doi: 10.1038/s41586-020-2342-5

27. Hong Y-K, Wu H-T, Ma T, Liu W-J, He X-J. Effects of Glycyrrhiza glabra polysaccharides on immune and antioxidant activities in high-fat mice. International Journal of Biological Macromolecules (2009) 45:61–64. doi: 10.1016/j.ijbiomac.2009.04.001

28. Ayeka PA, Bian Y, Githaiga PM, Zhao Y. The immunomodulatory activities of licorice polysaccharides (Glycyrrhiza uralensis Fisch.) in CT 26 tumor-bearing mice. BMC Complement Altern Med (2017) 17:536. doi: 10.1186/s12906-017-2030-7

29. Dexamethasone in Hospitalized Patients with Covid-19. New England Journal of Medicine (2021) 384:693–704. doi: 10.1056/NEJMoa2021436

30. Maddah M, Bahramsoltani R, Yekta NH, Rahimi R, Aliabadi R, Pourfath M. Proposing high-affinity inhibitors from Glycyrrhiza glabra L. against SARS-CoV-2 infection: virtual screening and computational analysis. New J Chem (2021) 45:15977–15995. doi: 10.1039/D1NJ02031E

31. Sinha SK, Prasad SK, Islam MA, Gurav SS, Patil RB, AlFaris NA, Aldayel TS, AlKehayez NM, Wabaidur SM, Shakya A. Identification of bioactive compounds from Glycyrrhiza glabra as possible inhibitor of SARS-CoV-2 spike glycoprotein and non-structural protein-15: a pharmacoinformatics study. Journal of Biomolecular Structure and Dynamics (2021) 39:4686–4700. doi: 10.1080/07391102.2020.1779132

32. Murck H. Symptomatic Protective Action of Glycyrrhizin (Licorice) in COVID-19 Infection? Frontiers in Immunology (2020) 11:1239. doi: 10.3389/fimmu.2020.01239

33. Safa O, Hassani-Azad M, Farashahinejad M, Davoodian P, Dadvand H, Hassanipour S, Fathalipour M. Effects of Licorice on clinical symptoms and laboratory signs in moderately ill patients with pneumonia from COVID-19: A structured summary of a study protocol for a randomized controlled trial. Trials (2020) 21:790. doi: 10.1186/s13063-020-04706-3

34. Qin C, Zhou L, Hu Z, Zhang S, Yang S, Tao Y, Xie C, Ma K, Shang K, Wang W, et al. Dysregulation of immune response in patients with COVID-19 in Wuhan, China. Clin Infect Dis doi: 10.1093/cid/ciaa248

35. Polidoro RB, Hagan RS, de Santis Santiago R, Schmidt NW. Overview: Systemic Inflammatory Response Derived From Lung Injury Caused by SARS-CoV-2 Infection Explains Severe Outcomes in COVID-19. Frontiers in Immunology (2020) 11: https://www.frontiersin.org/article/10.3389/fimmu.2020.01626 [Accessed May 11, 2022]

36. Tay MZ, Poh CM, Rénia L, MacAry PA, Ng LFP. The trinity of COVID-19: immunity, inflammation and intervention. Nat Rev Immunol (2020) 20:363–374. doi: 10.1038/s41577-020-0311-8

37. Shi Y, Wang Y, Shao C, Huang J, Gan J, Huang X, Bucci E, Piacentini M, Ippolito G, Melino G. COVID-19 infection: the perspectives on immune responses. Cell Death & Differentiation (2020) 27:1451–1454. doi: 10.1038/s41418-020-0530-3

38. Rizvi ZA, Tripathy MR, Sharma N, Goswami S, Srikanth N, Sastry JLN, Mani S, Surjit M, Awasthi A, Dikshit M. Effect of Prophylactic Use of Intranasal Oil Formulations in the Hamster Model of COVID-19. Frontiers in Pharmacology (2021) 12: https://www.frontiersin.org/article/10.3389/fphar.2021.746729 [Accessed May 11, 2022]

39. Bonifacius A, Tischer-Zimmermann S, Dragon AC, Gussarow D, Vogel A, Krettek U, Gödecke N, Yilmaz M, Kraft ARM, Hoeper MM, et al. COVID-19 immune signatures reveal stable antiviral T cell function despite declining humoral responses. Immunity (2021) 54:340–354.e6. doi: 10.1016/j.immuni.2021.01.008

40. Radermecker C, Detrembleur N, Guiot J, Cavalier E, Henket M, d’Emal C, Vanwinge C, Cataldo D, Oury C, Delvenne P, et al. Neutrophil extracellular traps infiltrate the lung airway, interstitial, and vascular compartments in severe COVID-19. Journal of Experimental Medicine (2020) 217:e20201012. doi: 10.1084/jem.20201012

41. Cheng O, Palaniyar N. NET balancing: a problem in inflammatory lung diseases. Frontiers in Immunology (2013) 4: https://www.frontiersin.org/article/10.3389/fimmu.2013.00001 [Accessed May 11, 2022]

42. Mlcochova P, Kemp SA, Dhar MS, Papa G, Meng B, Ferreira IATM, Datir R, Collier DA, Albecka A, Singh S, et al. SARS-CoV-2 B.1.617.2 Delta variant replication and immune evasion. Nature (2021) 599:114–119. doi: 10.1038/s41586-021-03944-y

43. McCallum M, Czudnochowski N, Rosen LE, Zepeda SK, Bowen JE, Walls AC, Hauser K, Joshi A, Stewart C, Dillen JR, et al. Structural basis of SARS-CoV-2 Omicron immune evasion and receptor engagement. Science (2022) 375:864–868. doi: 10.1126/science.abn8652

44. Girija PLT, Sivan N. Ayurvedic treatment of COVID-19/SARS-CoV-2: A case report. Journal of Ayurveda and Integrative Medicine (2020) doi: 10.1016/j.jaim.2020.06.001

45. Gomaa AA, Abdel-Wadood YA. The potential of glycyrrhizin and licorice extract in combating COVID-19 and associated conditions. Phytomedicine Plus (2021) 1:100043. doi: 10.1016/j.phyplu.2021.100043

46. Gamaleldin MMA. Impact of Different Treatment Modalities on Immunity Against COVID-19. [Clinical trial registration]. clinicaltrials.gov (2020). https://clinicaltrials.gov/ct2/show/NCT04553705 [Accessed May 9, 2022]

47. Cheel J, Antwerpen PV, Tůmová L, Onofre G, Vokurková D, Zouaoui-Boudjeltia K, Vanhaeverbeek M, Nève J. Free radical-scavenging, antioxidant and immunostimulating effects of a licorice infusion (Glycyrrhiza glabra L.). Food Chemistry (2010) 122:508–517. doi: 10.1016/j.foodchem.2010.02.060

48. Bordbar N, Karimi MH, Amirghofran Z. The effect of glycyrrhizin on maturation and T cell stimulating activity of dendritic cells. Cellular Immunology (2012) 280:44–49. doi: 10.1016/j.cellimm.2012.11.013

49. Chen Z, John Wherry E. T cell responses in patients with COVID-19. Nat Rev Immunol (2020) 20:529–536. doi: 10.1038/s41577-020-0402-6

50. Makni-Maalej K, Boussetta T, Hurtado-Nedelec M, Belambri SA, Gougerot-Pocidalo M-A, El-Benna J. The TLR7/8 Agonist CL097 Primes N-Formyl-Methionyl-Leucyl-Phenylalanine–Stimulated NADPH Oxidase Activation in Human Neutrophils: Critical Role of p47phox Phosphorylation and the Proline Isomerase Pin1. The Journal of Immunology (2012) 189:4657–4665. doi: 10.4049/jimmunol.1201007

51. Janke M, Poth J, Wimmenauer V, Giese T, Coch C, Barchet W, Schlee M, Hartmann G. Selective and direct activation of human neutrophils but not eosinophils by Toll-like receptor 8. J Allergy Clin Immunol (2009) 123:1026–1033. doi: 10.1016/j.jaci.2009.02.015

52. Hayashi F, Means TK, Luster AD. Toll-like receptors stimulate human neutrophil function. Blood (2003) 102:2660–2669. doi: 10.1182/blood-2003-04-1078

53. Hoarau C, Gérard B, Lescanne E, Henry D, François S, Lacapère J-J, Benna JE, Dang PM-C, Grandchamp B, Lebranchu Y, et al. TLR9 Activation Induces Normal Neutrophil Responses in a Child with IRAK-4 Deficiency: Involvement of the Direct PI3K Pathway. The Journal of Immunology (2007) 179:4754–4765. doi: 10.4049/jimmunol.179.7.4754

54. François S, Benna JE, Dang PMC, Pedruzzi E, Gougerot-Pocidalo M-A, Elbim C. Inhibition of Neutrophil Apoptosis by TLR Agonists in Whole Blood: Involvement of the Phosphoinositide 3-Kinase/Akt and NF-κB Signaling Pathways, Leading to Increased Levels of Mcl-1, A1, and Phosphorylated Bad. The Journal of Immunology (2005) 174:3633–3642. doi: 10.4049/jimmunol.174.6.3633

55. Nagase H, Okugawa S, Ota Y, Yamaguchi M, Tomizawa H, Matsushima K, Ohta K, Yamamoto K, Hirai K. Expression and Function of Toll-Like Receptors in Eosinophils: Activation by Toll-Like Receptor 7 Ligand. The Journal of Immunology (2003) 171:3977– 3982. doi: 10.4049/jimmunol.171.8.3977

56. Awasthi D, Nagarkoti S, Kumar A, Dubey M, Singh AK, Pathak P, Chandra T, Barthwal MK, Dikshit M. Oxidized LDL induced extracellular trap formation in human neutrophils via TLR-PKC-IRAK-MAPK and NADPH-oxidase activation. Free Radical Biology and Medicine (2016) 93:190–203. doi: 10.1016/j.freeradbiomed.2016.01.004

57. Costa MF, Jesus TI, Lopes BRP, Angolini CFF, Montagnolli A, Gomes L de P, Pereira GS, Ruiz ALTG, Carvalho JE, Eberlin MN, et al. Eugenia aurata and Eugenia punicifolia HBK inhibit inflammatory response by reducing neutrophil adhesion, degranulation and NET release. BMC Complementary and Alternative Medicine (2016) 16:403. doi: 10.1186/s12906-016-1375-7

58. Tao L, Xu M, Dai X, Ni T, Li D, Jin F, Wang H, Tao L, Pan B, Woodgett JR, et al. Polypharmacological Profiles Underlying the Antitumor Property of Salvia miltiorrhiza Root (Danshen) Interfering with NOX-Dependent Neutrophil Extracellular Traps. Oxid Med Cell Longev (2018) 2018:4908328. doi: 10.1155/2018/4908328

59. Kumar S, Dikshit M. Metabolic Insight of Neutrophils in Health and Disease. Frontiers in Immunology (2019) 10: https://www.frontiersin.org/article/10.3389/fimmu.2019.02099 [Accessed May 11, 2022]

60. Parker H, Albrett AM, Kettle AJ, Winterbourn CC. Myeloperoxidase associated with neutrophil extracellular traps is active and mediates bacterial killing in the presence of hydrogen peroxide. J Leukoc Biol (2012) 91:369–376. doi: 10.1189/jlb.0711387

61. Douda DN, Khan MA, Grasemann H, Palaniyar N. SK3 channel and mitochondrial ROS mediate NADPH oxidase-independent NETosis induced by calcium influx. Proceedings of the National Academy of Sciences (2015) 112:2817–2822. doi: 10.1073/pnas.1414055112

62. Vorobjeva NV, Chernyak BV. NETosis: Molecular Mechanisms, Role in Physiology and Pathology. Biochemistry Moscow (2020) 85:1178–1190. doi: 10.1134/S0006297920100065

63. Dikalov SI, Kirilyuk IA, Voinov M, Grigor’ev IA. EPR detection of cellular and mitochondrial superoxide using cyclic hydroxylamines. Free Radical Research (2011) 45:417–430. doi: 10.3109/10715762.2010.540242

64. Fortner KA, Blanco LP, Buskiewicz I, Huang N, Gibson PC, Cook DL, Pedersen HL, Yuen PST, Murphy MP, Perl A, et al. Targeting mitochondrial oxidative stress with MitoQ reduces NET formation and kidney disease in lupus-prone MRL-lpr mice. Lupus Science & Medicine (2020) 7:e000387. doi: 10.1136/lupus-2020-000387

65. Zimmermann M, Aguilera FB, Castellucci M, Rossato M, Costa S, Lunardi C, Ostuni R, Girolomoni G, Natoli G, Bazzoni F, et al. Chromatin remodelling and autocrine TNFα are required for optimal interleukin-6 expression in activated human neutrophils. Nat Commun (2015) 6:6061. doi: 10.1038/ncomms7061

66. Singh A, Singh V, Tiwari RL, Chandra T, Kumar A, Dikshit M, Barthwal MK. The IRAK-ERK-p67phox-Nox-2 axis mediates TLR4, 2-induced ROS production for IL-1β transcription and processing in monocytes. Cell Mol Immunol (2016) 13:745–763. doi: 10.1038/cmi.2015.62

67. Schroder K, Tschopp J. The inflammasomes. Cell (2010) 140:821–832. doi: 10.1016/j.cell.2010.01.040

68. Zhou R, Tardivel A, Thorens B, Choi I, Tschopp J. Thioredoxin-interacting protein links oxidative stress to inflammasome activation. Nat Immunol (2010) 11:136–140. doi: 10.1038/ni.1831

69. Guo H, Callaway JB, Ting JP-Y. Inflammasomes: mechanism of action, role in disease, and therapeutics. Nat Med (2015) 21:677–687. doi: 10.1038/nm.3893

70. Münzer P, Negro R, Fukui S, di Meglio L, Aymonnier K, Chu L, Cherpokova D, Gutch S, Sorvillo N, Shi L, et al. NLRP3 Inflammasome Assembly in Neutrophils Is Supported by PAD4 and Promotes NETosis Under Sterile Conditions. Frontiers in Immunology (2021) 12: https://www.frontiersin.org/article/10.3389/fimmu.2021.683803 [Accessed May 11, 2022]

71. Nagarkoti S, Sadaf S, Awasthi D, Chandra T, Jagavelu K, Kumar S, Dikshit M. L-Arginine and tetrahydrobiopterin supported nitric oxide production is crucial for the microbicidal activity of neutrophils. Free Radical Research (2019) 53:281–292. doi: 10.1080/10715762.2019.1566605

72. Shukla S, Mehta A, John J, Mehta P, Vyas SP, Shukla S. Immunomodulatory activities of the ethanolic extract of Caesalpinia bonducella seeds. J Ethnopharmacol (2009) 125:252–256. doi: 10.1016/j.jep.2009.07.002

73. Solanki YB, Jain SM. Immunostimolatory activities of Vigna mungo L. extract in male Sprague–Dawley rats. Journal of Immunotoxicology (2010) 7:213–218. doi: 10.3109/15476911003792278

74. Yin C, Wu C, Du X, Fang Y, Pu J, Wu J, Tang L, Zhao W, Weng Y, Guo X, et al. PRL2 Controls Phagocyte Bactericidal Activity by Sensing and Regulating ROS. Frontiers in Immunology (2018) 9: https://www.frontiersin.org/article/10.3389/fimmu.2018.02609 [Accessed May 11, 2022]

75. Jyoti A, Singh AK, Dubey M, Kumar S, Saluja R, Keshari RS, Verma A, Chandra T, Kumar A, Bajpai VK, et al. Interaction of Inducible Nitric Oxide Synthase with Rac2 Regulates Reactive Oxygen and Nitrogen Species Generation in the Human Neutrophil Phagosomes: Implication in Microbial Killing. Antioxidants & Redox Signaling (2014) 20:417–431. doi: 10.1089/ars.2012.4970

76. Rizvi ZA, Puri N, Saxena RK. Evidence of CD1d pathway of lipid antigen presentation in mouse primary lung epithelial cells and its up-regulation upon Mycobacterium bovis BCG infection. PLOS ONE (2018) 13:e0210116. doi: 10.1371/journal.pone.0210116

77. Rizvi ZA, Dalal R, Sadhu S, Kumar Y, Kumar S, Gupta SK, Tripathy MR, Rathore DK, Awasthi A. High-salt diet mediates interplay between NK cells and gut microbiota to induce potent tumor immunity. Sci Adv (2021) 7:eabg5016. doi: 10.1126/sciadv.abg5016

78. Roy S, Rizvi ZA, Clarke AJ, Macdonald F, Pandey A, Zaiss DMW, Simon AK, Awasthi A. EGFR-HIF1α signaling positively regulates the differentiation of IL-9 producing T helper cells. Nat Commun (2021) 12:3182. doi: 10.1038/s41467-021-23042-x

79. Malik S, Sadhu S, Elesela S, Pandey RP, Chawla AS, Sharma D, Panda L, Rathore D, Ghosh B, Ahuja V, et al. Transcription factor Foxo1 is essential for IL-9 induction in T helper cells. Nat Commun (2017) 8:815. doi: 10.1038/s41467-017-00674-6

80. Nagarkoti S, Dubey M, Awasthi D, Kumar V, Chandra T, Kumar S, Dikshit M. S-Glutathionylation of p47phox sustains superoxide generation in activated neutrophils. Biochim Biophys Acta Mol Cell Res (2018) 1865:444–454. doi: 10.1016/j.bbamcr.2017.11.014

81. Singh AK, Awasthi D, Dubey M, Nagarkoti S, Kumar A, Chandra T, Barthwal MK, Tripathi AK, Dikshit M. High oxidative stress adversely affects NFκB mediated induction of inducible nitric oxide synthase in human neutrophils: Implications in chronic myeloid leukemia. Nitric Oxide (2016) 58:28–41. doi: 10.1016/j.niox.2016.06.002

